# Non-uniform distribution of myosin-mediated forces governs red blood cell membrane curvature through tension modulation

**DOI:** 10.1101/668582

**Authors:** Haleh Alimohamadi, Alyson S. Smith, Roberta B. Nowak, Velia M. Fowler, Padmini Rangamani

## Abstract

The biconcave disk shape of the mammalian red blood cell (RBC) is unique to the RBC and is vital for its circulatory function. Due to the absence of a transcellular cytoskeleton, RBC shape is determined by the membrane skeleton, a network of actin filaments cross-linked by spectrin and attached to membrane proteins. While the physical properties of a uniformly distributed actin network interacting with the lipid bilayer membrane have been assumed to control RBC shape, recent experiments reveal that RBC biconcave shape also depends on the contractile activity of nonmuscle myosin IIA (NMIIA) motor proteins. Here, we use the classical Helfrich-Canham model for the RBC membrane to test the role of heterogeneous force distributions along the membrane and mimic the contractile activity of sparsely distributed NMIIA filaments. By incorporating this additional contribution to the Helfrich-Canham energy, we find that the RBC biconcave shape depends on the ratio of forces per unit volume in the dimple and rim regions of the RBC. Experimental measurements of NMIIA densities at the dimple and rim validate our prediction that (a) membrane forces must be non-uniform along the RBC membrane and (b) the force density must be larger in the dimple than the rim to produce the observed membrane curvatures. Furthermore, we predict that RBC membrane tension and the orientation of the applied forces play important roles in regulating this force-shape landscape. Our findings of heterogeneous force distributions on the plasma membrane for RBC shape maintenance may also have implications for shape maintenance in different cell types.

## 1 Introduction

Cell shape and function are intricately coupled; cells must maintain specific shapes to migrate, divide normally, form tissues and organs during development, and support their physiological functions [1,2]. Maintenance or modification of cell shape is a concerted action of the actomyosin network at the whole cell level that allows for a stable actin network in polarized cells or a rapidly remodeling actin network for cell spreading and motility [3,4]. Thus, networks of actin filaments (F-actin) and the F-actin-activated motor protein non-muscle myosin II (NMII) specify cell shape by exerting forces on the plasma membrane to control membrane tension and curvature [5–8]. These actomyosin networks determine cell shapes and interactions during tissue morphogenesis in development [6,9–13] and their dysregulation has been implicated in cancer, [14–16], hearing disorders [17], podocyte filtration in the kidney [17,18], and neurodegeneration [19], among other physical issues. Local, nanoscale changes in actomyosin organization can lead to micron-scale changes in membrane curvature and cell shape to support normal cell function [20].

Human red blood cells (RBCs) have a biconcave disk shape, with a thin central dimple region surrounded by a thicker rim [21,22] (Fig. 1). This shape enables efficient gas and ion exchange and increases RBC deformability and resiliency in the circulation [23–25]. Deviations from biconcavity interfere with RBC function in diseases such as congenital hemolytic anemias [23,24,26], sickle cell disease [27], and malaria [28,29]. Due to its lack of transcellular cytoskeleton or internal organelles, RBC shape depends exclusively on the plasma membrane, and has long served as a simple model for membrane structure and function [30]. The RBC membrane is supported by the membrane skeleton, a two-dimensional network of short F-actins interconnected by long, flexible spectrin molecules [30,31], which bind to transmembrane proteins to maintain membrane tension, curvature, and mechanical properties of the RBC [23,24,30,32,33].

**Figure 1:**
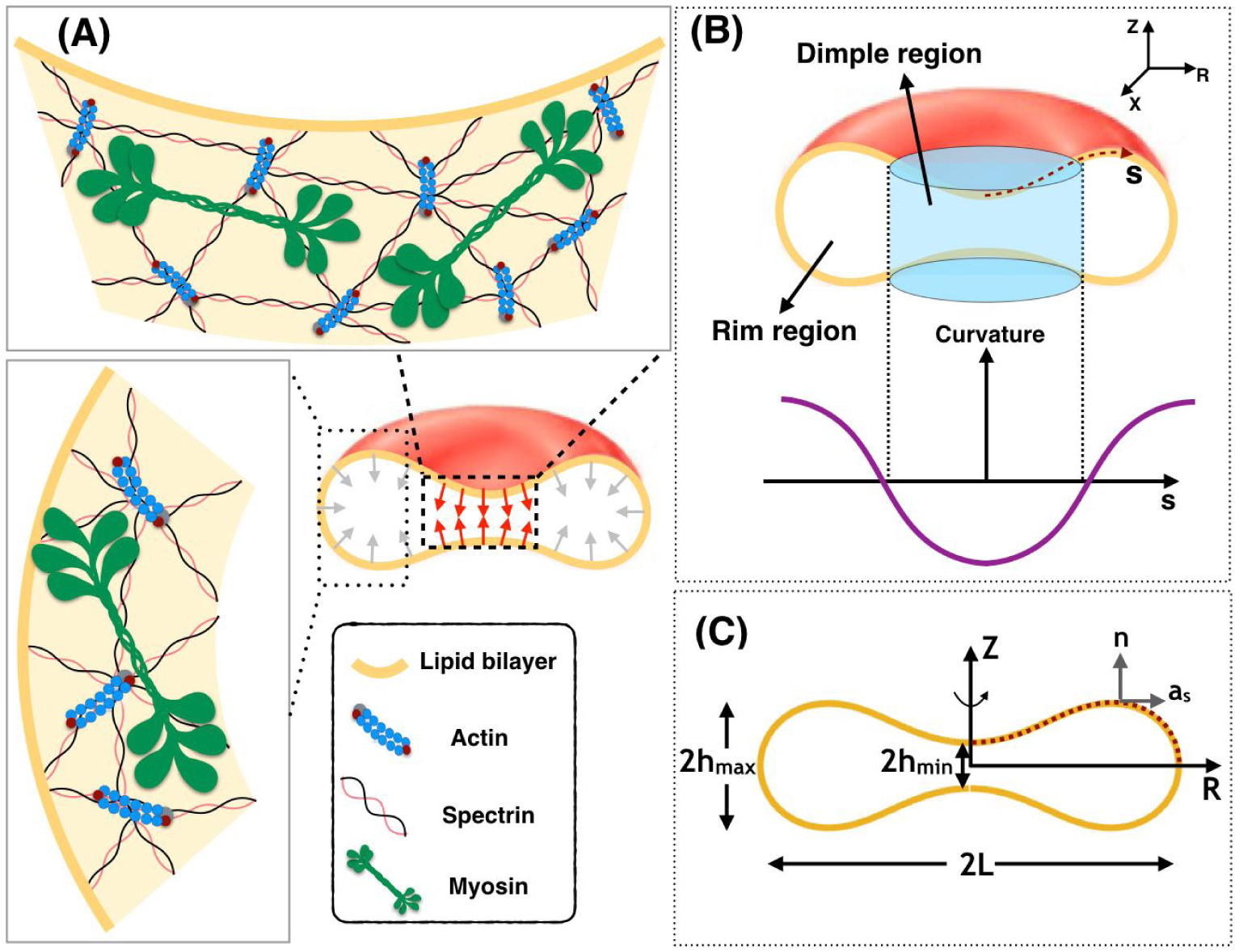
Interaction of the membrane and skeleton controls the shape of the RBC. (A) Schematic depiction of the biconcave disk shape of an RBC plasma membrane and the membrane skeleton underneath. The effect of NMIIA filaments (shown in green) is modeled by local forces applied to the plasma membrane (red and gray arrows). (B) Two distinct regions are identified in a biconcave RBC -- the dimple and the rim regions. In the dimple region (blue cylinder), each RZ cross-section of the shape has a negative curvature along its arclength. In contrast, at the rim, the curvature of each RZ section is positive along the arclength. (C) The geometry of a simulated RBC in axisymmetric coordinates and the three characteristic length scales that represent the biconcave shape of the RBC. 2h_min_ is the minimum height at the dimple, 2h_max_ is the maximum height at the rim, and 2L denotes the cell’s maximum diameter. The dotted red curve shows the computational domain for our mechanical model. **n** is the unit normal vector to the membrane surface and **a**_s_ is the unit tangent vector in the direction of arclength.

Historically, the biconcave disk shape of the RBC has been modeled as a thin elastic shell using the Helfrich-Canham energy model, treating the RBC as a lipid bilayer whose properties are spatially homogeneous along the entire RBC membrane [34,35]. This model, a classic in the field of membrane mechanics, was able to explain the observed RBC shape as a family of solutions for a given area and volume [34,35]. Subsequent extensions of this model include consideration of reduced volume [34,35], leaflet asymmetry (also termed spontaneous curvature) [36], and lateral distribution of membrane constituents [37–39]. Additional refinements have modeled the membrane as a two-component system comprised of an incompressible lipid bilayer associated with an elastic spectrin-actin network uniformly distributed along the membrane [40–42].

To date, computational models to account for the biconcave disk shape of the RBC have not considered the contribution of actomyosin contractility, yet RBCs contain NMII. NMIIA is the predominantly expressed isoform and has biochemical properties similar to NMIIs in other cell types [43,44]. Due to its low abundance, a potential role for NMIIA in RBC shape had been largely ignored by experimental biologists. However, we showed recently in Smith et al. [45] that RBC NMIIA forms bipolar filaments that bind to the membrane skeleton F-actin via their motor domains to control RBC membrane tension, biconcave disk shape and deformability [45] (Fig. 1A). Specifically, blebbistatin inhibition of NMIIA motor activity in RBCs leads to loss of the biconcavity and the formation of elongated shapes, indicating an important role for NMIIA-generated forces in maintaining RBC biconcave disk shape and deformability [45]. Notably, the NMIIA filaments are sparsely distributed along the RBC membrane (∼0.5 filaments per square micrometer), and thus would be expected to apply localized forces to the membrane and skeleton [45]. However, it remains unclear how the magnitude and distribution of NMII-mediated localized forces could provide a mechanism to influence membrane curvature with respect to the morphology of RBCs.

In this study, motivated by our recent experimental observation [45], we investigated the role of local forces in modulating the shape of the RBC. We revisit the classical Helfrich-Canham model and modify it to account for localized forces representing the NMIIA-generated forces on the plasma membrane. By adding this extra degree of freedom to the classical Helfrich-Canham model, we sought to focus on how forces applied to the membrane, rather than spontaneous curvature or reduced volume, can result in the shapes that are comparable to the experimentally observed RBC shapes. To determine the set of force distributions that most closely reproduce experimentally observed RBC shapes, we varied the applied force heterogeneously along the membrane. Our model predicts that the best match between simulations and experiments for RBC shapes is obtained when there are two force-dependent curvature domains -- a dimple region with negative curvature and a rim with positive curvature and lower forces at the cell periphery (Fig. 1B). The dimple region needs greater forces and the rim lower forces, and this force heterogeneity results in the experimentally observed curvature.

Experimental measurements of the NMIIA distribution validated our prediction of non-uniform force distribution; more NMIIA puncta are found in the dimple region than in the rim. Our model also predicts that membrane tension and the orientation of the applied forces are the important parameters that regulate RBC morphology and the force density distribution across the dimple and rim regions of a biconcave RBC. These findings can provide a potential design principle for RBC shape maintenance despite variations in NMIIA puncta localizations. Additionally, membrane skeleton spectrin-F-actin networks, first discovered in RBCs, are present at the plasma membrane of all metazoan cells [46–48]. Our computational modeling of the RBC biconcave disk shape provides a simple system to elucidate how actomyosin contractility can control cell shape via micron-scale localized forces on the membrane associated skeleton.

## 2 Model development

The RBC membrane is a thin elastic material that can bend but resists stretching. This feature enables the RBC to deform and adjust its shape in response to applied stresses. Here, we outline the governing equations of our model. This approach will allow us to predict how the induced surface forces by NMIIA motor activity can regulate the biconcave morphology of RBCs in mechanical equilibrium. To develop our model and solve it numerically, we make the following assumptions.

### 2.1 Assumptions

1. We consider that the radii of membrane curvatures are much larger than the thickness of the bilayer [36]. This allows us to treat the lipid bilayer as a thin elastic shell and model the bending energy of the membrane by the Helfrich–Canham energy, which depends only on the local curvatures of the surface and compositional heterogeneities [34,35].
2. Due to the high stretching modulus of lipid bilayers, we assume that the membrane is locally incompressible [49]. We use a Lagrange multiplier to implement this constraint [50,51].
3. We assume that the RBC is at mechanical equilibrium at all time scales, allowing us to neglect inertia [52–54]. This assumption is consistent with the experimentally observed shapes for the resting RBCs in both *vivo* and *vitro* [55,56].
4. We assume that the total surface area of the RBC membrane is constant (∼135 *μ*m^2^) [57] [58].
5. For simplicity in the numerical simulations, we assume that the RBC is rotationally symmetric and also has a reflection symmetry with respect to the Z = 0 plane (see Fig. 1C) [34,57,59,60]. This assumption reduces the computational cost of the simulation to simply calculating the shape of the curve shown by the red dotted line in Fig. 1C.

### 2.2 Membrane mechanics

In mechanical equilibrium, the shape of the membrane in response to an applied force can be obtained as a result of minimization of the membrane bending energy and the work done by the applied forces by the cytoskeleton. The total energy in this case is given by [61,62]

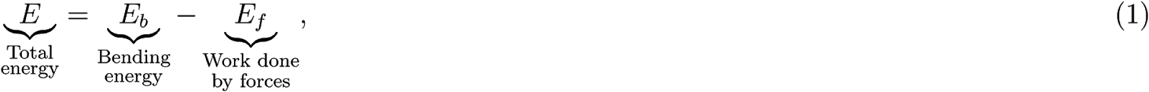

where is *E* the total energy of the system, *E*_*b*_ is the bending energy and *E*_*f*_ is the work done by the applied forces given by

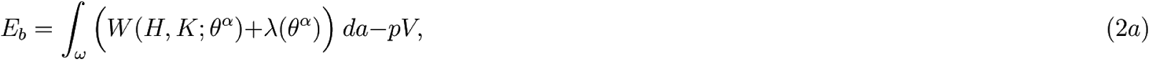

and

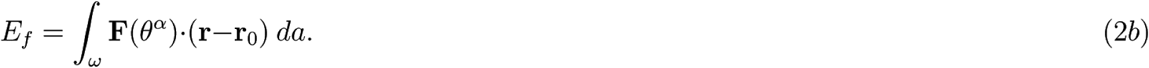

Here, *ω* is the total surface area of the membrane, *W* is the energy density per unit area *θ^α^*, denotes the surface coordinate where *α* ∈ {1,2}, *H* is the mean curvature, *K* is the Gaussian curvature, *λ* is the membrane tension, *p* is the pressure difference across the membrane, *V* is the enclosed volume, **F** is the force density per unit area representing the applied force density to the membrane surface by the NMIIA motor proteins, r is the position vector in the current configuration, and r_0_ is the position vector in the reference configuration. The bending energy of the membrane is modeled using the Helfrich-Canham energy, defined by [34,35,50,63,64],

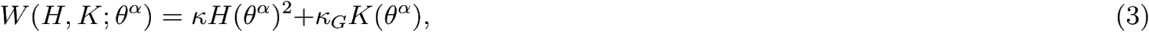

where *κ* and *κ*_*G*_ are constants representing the bending and Gaussian moduli respectively [65]. To minimize the bending energy (Eq.1) and obtain the RBC shapes from simulations under the action of local forces, we used the variational approach which yields the so-called “shape equation” [63,66],

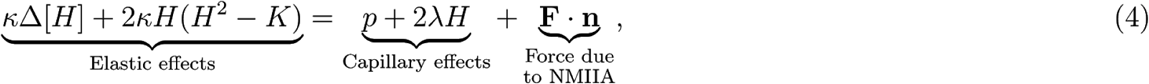

where Δ is the surface Laplacian (also known as the Beltrami operator).

The incompressibility condition for the lipid membrane results in the spatial variation of membrane tension given by [63,66]

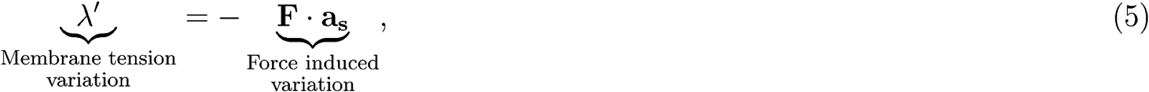

where (·)′ is the derivative with respect to the arclength. The shape equation (Eq. 4) along with the incompressibility condition (Eq. 5) represents the relationship between the forces applied by NMIIA motor proteins and the resulting shape of RBCs. A complete derivation of the governing equations of the force balance, the notations used, and the non-dimensionalization procedure are presented in the supplementary online material (SOM). We refer the interested reader to [50,61] for details on the mathematical principles underlying these models.

### 2.3 Parametrization of RBC biconcave morphology and shape error estimation

The geometry of human RBCs has been studied extensively using a variety of different methods such as light microscopy [67,68], interference holography [69,70], resistive pulse spectroscopy [71], micropipette aspiration [72,73], and light scattering [74,75]. In Fig. 2A, we summarize the reported values for the RBC geometrical parameters from the literature [67,69,72,75–77] in terms of the three characteristic lengths (h_min_, h_max_ and L) (Fig. 1C), the volume (V), the surface area (A), and the sphericity index (SI).

**Figure 2:**
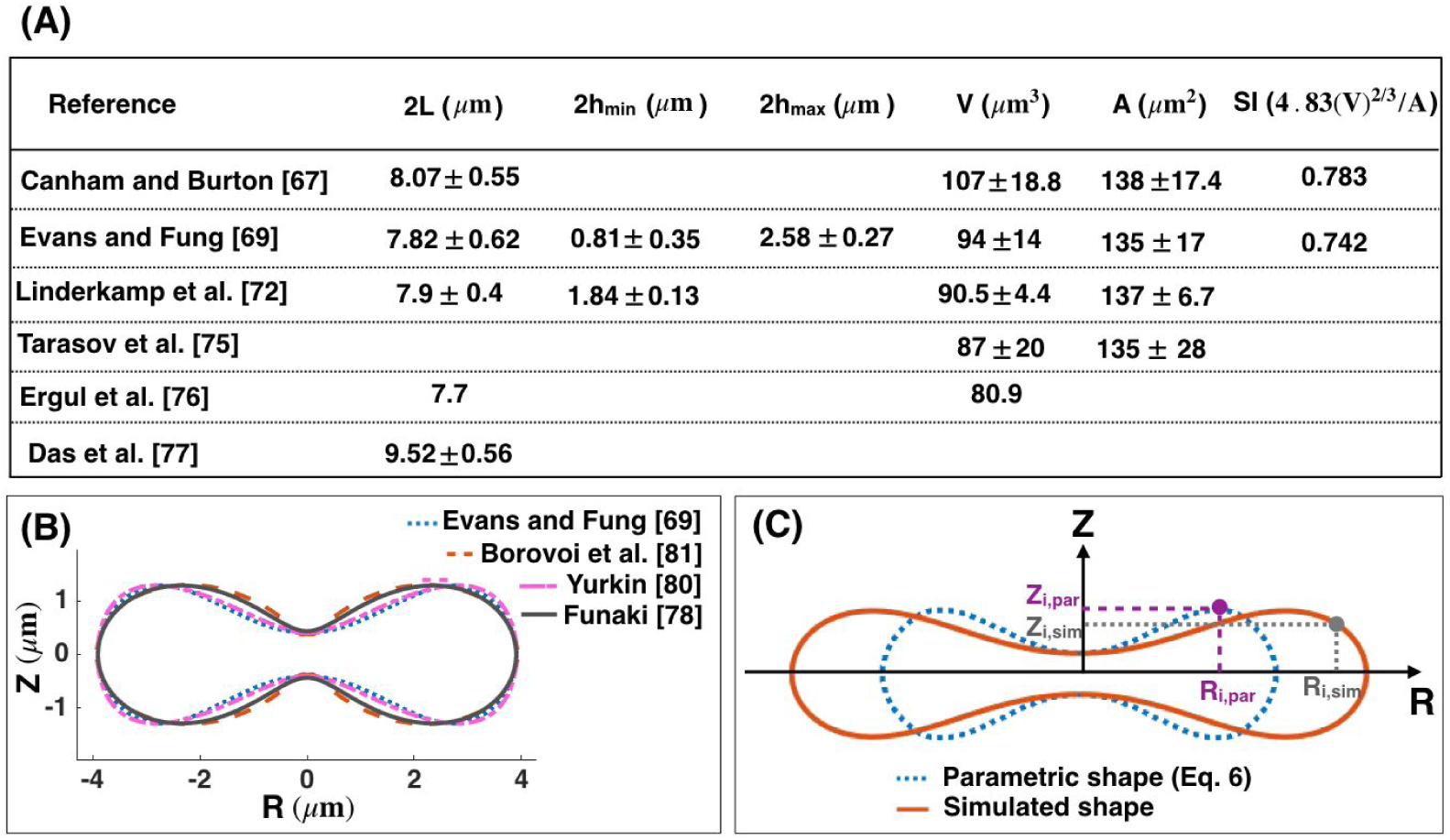
(A) Healthy human RBC dimensions from the literature [67,69,72,75–77]. (B) Comparison between the proposed parametric models describing the biconcave morphology of an RBC. There is a close match between the four models for the fixed minimum height of the dimple, maximum height of the rim, and the maximum diameter (C) Discretization scheme of the parametric shape of an RBC (Eq. 6) (dotted blue line) and the simulated geometry obtained from our mechanical model (Eqs. 4 and 5) (solid red line). Each experimental and simulated shape is discretized into *N* nodes where *i* indicates the node index. These nodes are used to compute the total error in the simulated RBC geometry (Eq. 8).

During the last few decades, several parametric models have been proposed to describe the biconcave morphology of the RBC [69,70,78–82]. Funaki proposed the Cassini oval model with two coefficients to represent the RBC geometry [78]. Kuchel et al. [79] and later Yurkin [80] modified the Cassini oval model to implicit equations with three and four coefficients, respectively. Borovoi et al. introduced a function in spherical coordinates to characterize the RBC morphology [81]. The most realistic model was proposed by Evans and Fung [69], where they first obtained images from 50 human RBC samples using light microscopy and then fitted a parametric equation to the RZ cross-sectional shape of the RBCs (Fig. 1C) using statistical analysis. The Evans and Fung proposed function is given by

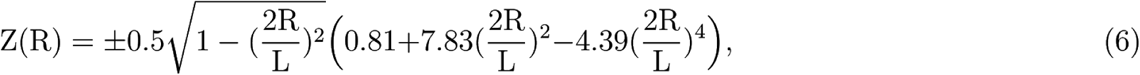

where *R* is the radius from the axis of rotation and *Z* is the height from the base plane. In Fig. 2B, we plotted the different proposed parametric models for the biconcave shape of an RBC. We observed that for the fixed height of the dimple (h_min_), height of the rim (h_max_), and the maximum diameter (L), all models generate similar shapes, but with slight differences. In this study, we used the Evans and Fung parametric equation in Eq. 6 as the reference data for the experimental shape of an RBC, since Eq. 6 was developed based on the direct experimental measurement and fit well with the observed RBC shapes [83,84].

To quantify the deviation between simulated geometries obtained from our mechanical model and the parametric shape equation for the RBC (Eq. 6), we define three errors ϵ_min,_ ϵ_max_ and ϵ_L_ as

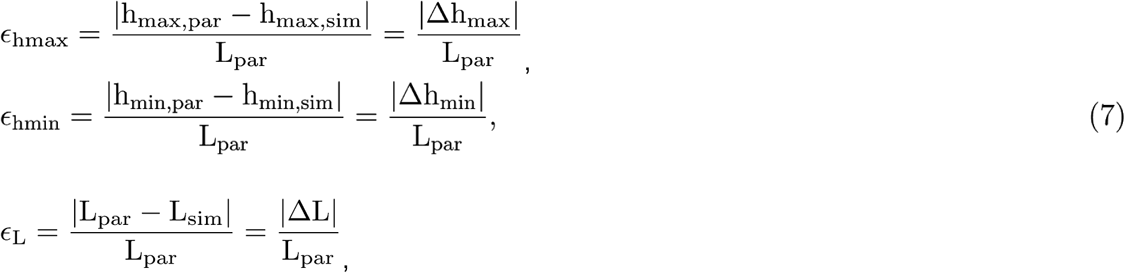

where (.)_,par_ is the experimentally measured length scale fitted to the parametric equations (Eq. 6) and (.)_,sim_ is the length scale obtained from the numerical simulation (Eqs. 4 and 5). The total error (*ϵ*_total_) in the shape of the simulated RBCs is then calculated by the root mean square between every two mapped points in the parametric shape of an RBC and the simulated geometries (Fig. 2C) given by

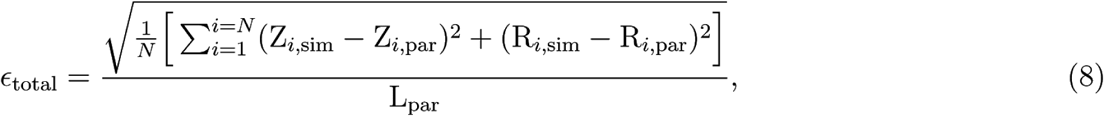

where *N* is the total number of nodes across the RBC shapes, *i* is the index node, Z_*i*,sim_ and Z_*i*,par_ are the height of the simulated and the parametric (Eq. 6) RBC shape at index *i*, respectively. R_*i*,sim_ is the radius of the simulated shape at index *i*, and R_*i*,par_ is the radius of the RBC parametric shape (Eq. 6) at index *i*.

While Eq. 8 represents the error in the simulated shapes compared to the RBC parametric shape, it does not capture measurement errors as Eq. 6 was developed based on the average dimensions of experimentally observed RBCs. However, there are standard deviations in the measured dimensions as reported by Evans and Fung [69] and resolution of imaging methods introduces additional uncertainties. Here, to account for these uncertainties, we assume that the given parametric equation by Evans and Fung [69] can be written as

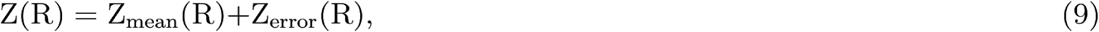

where Z_mean_(R) is the given function in Eq. 6 and we define Z_error_(R) as the fitting error of the Evans and Fung parametric equation to the actual shape of an RBC. In this study, we assume that Z_error_(R) is approximately 10% of Z_mean_(R) in order to represent the variance of RBC dimensions.

### 2.4 Numerical simulation

Simplifying the shape equation (Eqs. 4 and 5) for a rotationally symmetric RBC gives us a set of first order differential equations (Eq. S11). In order to obtain the RBC shapes from simulations and determine the role of NMIIA-generated forces in maintaining the biconcave morphology, we need to solve the coupled differential equations (Eq. S11) for the defined boundary conditions (Eq. S12). Here, we used the commercially available finite element solver COMSOL MULTIPHYSICS 5.3a to solve the governing differential equations (Eqs. S11 and S12). In all our simulations, the transmembrane pressure is set to zero (*p* =0).

## 3 Results

### 3.1 Uniform distribution of force density across the membrane surface is not sufficient to recover the biconcave shape of an RBC

Modeling studies of RBC shapes have been based on the assumption that the RBC membrane and skeleton are spatially homogeneous [23,24,30,65]. Therefore, we first performed simulations with a uniform pulling force density (F_uniform_) applied normally on the membrane surface (Figs. 3A, B). This uniform pulling force density can be interpreted as a pressure difference between the inside and outside of the RBC which specifies the change in the RBC volume compared to an equivalent sphere (the reduced volume) [36,85,86]. Considering the RBC membrane rigidity and elasticity, experimental measurements showed that a small pressure difference -- on order of ∼1 pN/*μ*m^2^-- is sufficient to form and maintain a biconcave RBC from a spherical cell [86–89].

**Figure 3:**
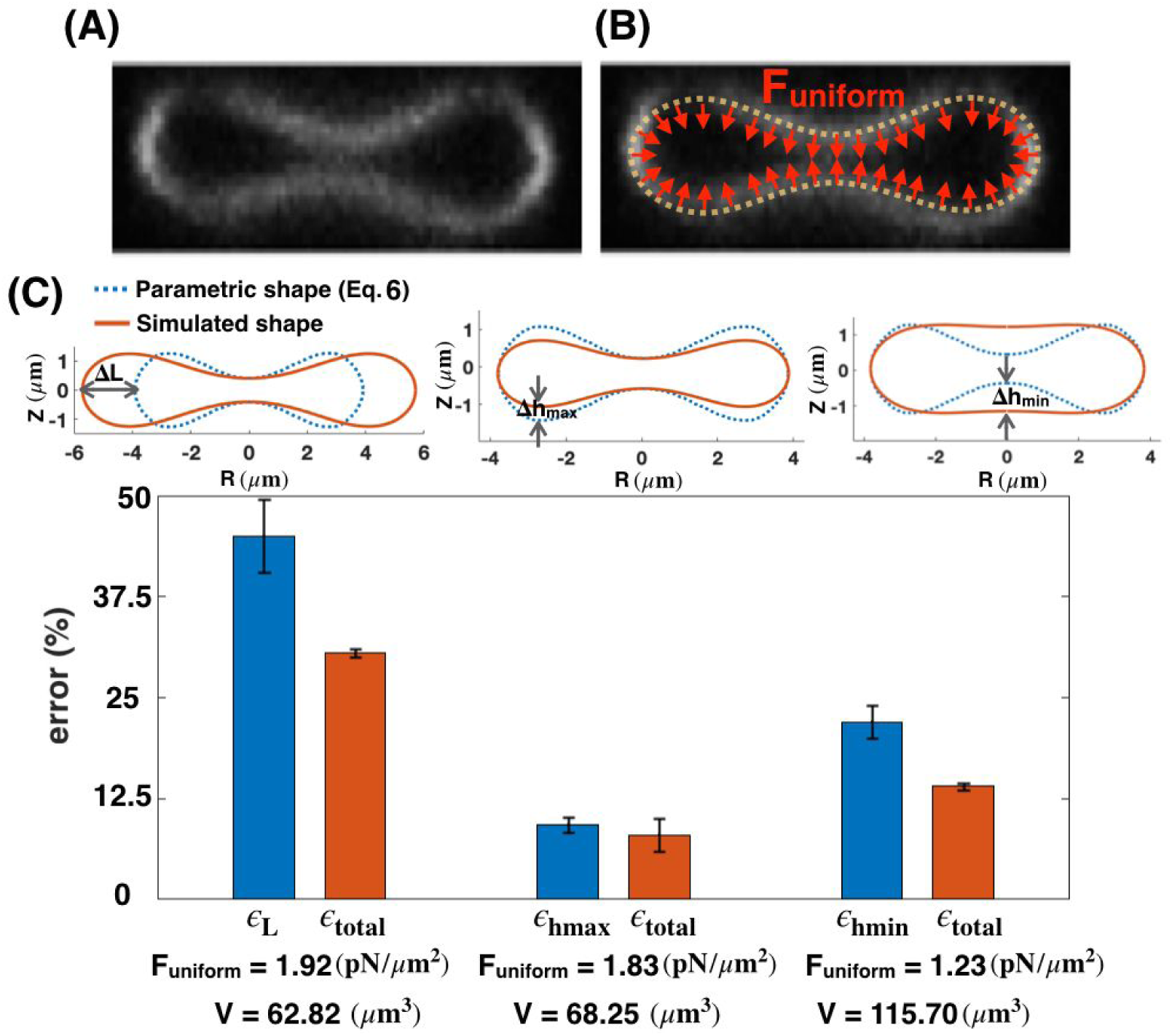
Mismatch between the parametric shape of an experimentally observed RBC (Eq. 6) and the shapes obtained from simulations (Eqs. S10) with a uniform distribution of the pulling force density across the membrane surface. (A) RZ view of the center of an RBC from a confocal Z-stack of an RBC stained for the membrane marker glycophorin A. (B) Schematic of a biconcave RBC with a uniform distribution of the normal pulling force density (red arrows). F_uniform_ represents the magnitude of the pulling force density. (C) Calculated error in the characteristic length scales (Eq. 7), and total shape error (Eq. 8) for different values of the force density. The total shape error (ϵ_total_) calculated by Eq. 8 is minimum for F_uniform_= 1.83 pN/*μ*m^2^, when there is only a mismatch in the maximum height of the RBC morphology (center bar). For all three values of the applied uniform force densities, the calculated volume (V) is shown on the X-axis and is significantly different from the reported experimental data.

To perform our simulations, we assumed that the RBC membrane area is large enough that the lateral membrane tension is negligible (*λ* = 0) [90–92]. We also set the bending modulus to be in the range of physiologically reported values for the RBC membrane (*κ*= 9 × 10^−19^ J) [91]. As shown in Fig. 3B, for a given value of uniform pulling force, we were able to match two out of three characteristic length scales of the simulated shapes with the parametric shape of an experimentally observed RBC (Eq. 6). Furthermore, we can see that for all configurations in Fig. 3C, the calculated uniform force density from our mechanical model is in the order of the reported pressure difference for a biconcave RBC (F_uniform_ ∼ O(1) pN/*μ*m^2^), which validates the accuracy of our numerical results (Fig. 3C, bottom row).

Based on the results in Fig. 3C, we observe that for the large value of the pulling force density (F_uniform_ = 1.92 pN/*μ*m^2^), the maximum and the minimum heights of the simulated shape match well with the parametric shape, while the maximum diameter does not (Fig. 3B left). For the intermediate pulling force density (F_uniform_ = 1.83 pN/*μ*m^2^), the minimum height and the maximum diameter of the simulated shape are in good agreement with the parametric shape, but the maximum height is not (Fig. 3C center). Finally, for the small pulling force density (F_uniform_= 1.23 pN/*μ*m^2^), the mismatch between the simulated geometry and the parametric shape of the RBC is in the minimum height of the dimple (Fig. 3C right).

For each value of the applied pulling force density, we plotted the error for each of the characteristic lengths (L, h_max_ or h_min_) (Eq. 7) and the total error (Eq. 8) (Fig. 3C). We found that both characteristic and the total shape errors have the lowest value (ϵ_hmax_∼ 10.23% and ϵ_total_∼ 8.2%) at the intermediate uniform force density, when there is only a relatively small mismatch in the maximum height (Δh_max_) (Fig. 3C center). Thus, we can predict that among the three main characteristic length scales of an RBC, the maximum height of the rim (2h_max_) appears to be the least critical dimension in order to minimize the shape error of the simulated geometries. It should be mentioned that for each case here, we first calculated the mean errors based on the given parametric equation (Eq. 6) and then we computed the error bars using Eq. 9.

In addition to the biconcave shape of an RBC, the volume of the RBC is one of the critical parameters that is regulated by multiple transport systems [86]. Thus, in Fig. 3C, we calculated the volume of each simulated geometry (V) using Eq. S13b (Fig. 3C). Based on our results, for all three values of the uniform force densities, the volume of the simulated shapes are far from the reported experimental data in Fig. 2A. For the large and the intermediate force densities (F_uniform_= 1.92 pN/*μ*m^2^ and F_uniform_= 1.83 pN/*μ*m^2^), the volumes of the simulated geometries (Fig. 3C) are much smaller than the reported values which range from V= 80 *μ*m^3^ to V= 107 *μ*m^3^ given in Fig. 2A. In contrast, for the small force density (F_uniform_ = 1.23 pN/*μ*m^2^), the volume of the shape obtained from the simulation is significantly larger than the experimental values (Fig. 3C).

### 3.2 Local force density at the RBC dimple reduces the shape error

Given that the shape mismatch and volume difference of the simulated RBC are relatively large compared to the experimentally measured RBC dimensions in Fig. 2, we next asked if we could change the distribution of the non-uniform pulling force density to reduce the shape error and obtain a better agreement between the experimentally reported shapes for RBCs and our model. We conducted simulations of Eqs (S10, S11) but assuming that the applied normal force per unit area is locally concentrated in the dimple region (F_dimple_) and that there is no force along the surface of the rim (Figs. 4A, B). This heterogeneous force distribution along the membrane was implemented using a hyperbolic tangent function (Eq. S23).

**Figure 4:**
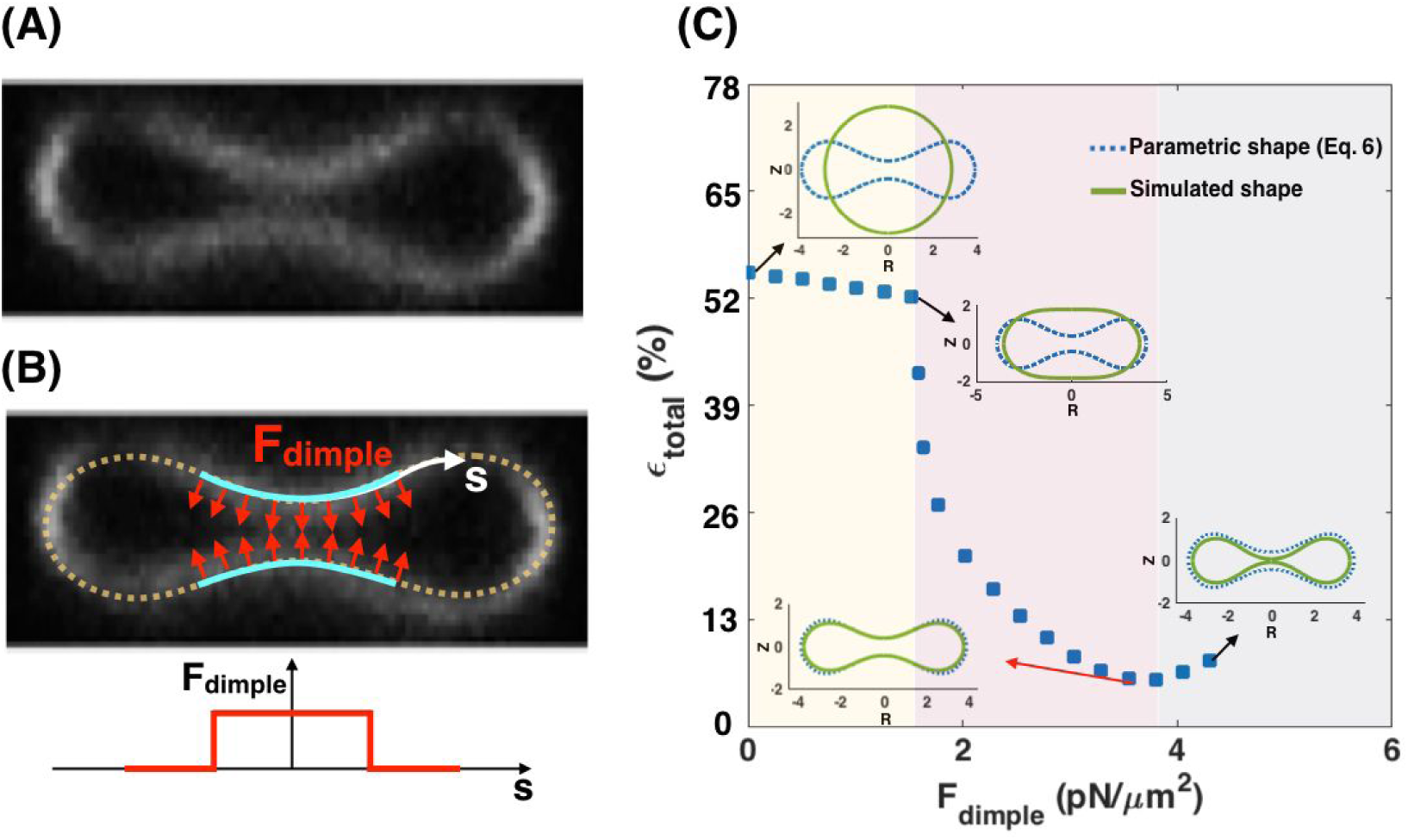
A local distribution of the pulling force density at the RBC dimple results in a better match between the parametric shape of an experimental observed RBC (Eq. 6) and the shape obtained from the simulation. (A) RZ view of the center of an RBC from a confocal Z-stack of an RBC stained for the membrane marker glycophorin A. (B, upper) A schematic depicting a biconcave RBC with a local force at the dimple area (red arrows) and no force in the rim region. F_dimple_ represents the magnitude of the pulling force density in the dimple region. (B, lower) The applied force density at the dimple as a function of the arclength (Eq. S23). (C) The simulated shape of the RBC with a local pulling force density in the dimple (solid green line) in comparison with the RBC parametric shape (dotted blue line). (C) The nonlinear behavior of the total error when increasing the dimple force density (F_dimple_). Three different regimes can be identified based on the shape of the simulated RBC; *(i)* the spherical shapes where h_max_ = h_min_ for the low F_dimple_ (yellow area), *(ii)* the biconcave shapes where the dimple forms (h_max_ > h_min_) for the mid-range of F_dimple_ (purple area), and *(iii)* the kissing shapes where h_min_ → 0 for large F_dimple_ (gray area). The shape error has the lowest value at F_dimple_ = 3.73 pN/*μ*m^2^ (ϵ _total_ ∼ 5.62%) when the minimum height of the dimple in the simulated geometry matches closely with the minimum height of the parametric shape. The volume of the simulated RBC at F_dimple_ = 3.73 pN/*μ*m^2^ is about 76.78 *μ*m^3^.

In Fig. 4C, we compare the RBC shapes obtained from the simulation with the application of increased local pulling force density at the dimple. We find that the total error is a nonlinear function of F_dimple_; as F_dimple_ increases, the total error in shape mismatch decreases and then increases again. Based on the shape of the simulated RBC, we can identify three different regimes (Fig. 4C). For low dimple force density (F_dimple_ < 1.81 pN/*μ*m^2^), the simulated geometry has a spherical shape (h_max_ = h_min_) and therefore the shape error is large (ϵ_total_ > 50%) (yellow area in Fig. 4C). With increasing the magnitude of dimple force density (1.81 pN/*μ*m2 < F_dimple_< 3.73 pN/*μ*m^2^), the dimple forms—biconcave shapes where h_max_ > h_min_ — and the shape error decreases sharply (purple area in Fig. 4C). When higher levels of force are applied at the dimple (F_dimple_ > 3.73 pN/*μ*m^2^), the error increases because the distance between the two bilayers in the dimple becomes too narrow (kissing shapes where hmin = 0) (Fig. 4C). We also observed a similar nonlinear trend in the calculated errors for the characteristic lengths (Eq. 7) as a function of dimple force density (Fig. S1).

Based on our results in Fig. 4C, the shape error has a minimum value of ϵ_total_ ∼ 5.62% for the case where F_dimple_ = 3.73 pN/*μ*m^2^. This total error is less than that for all the simulated shapes determined in the case of a uniform force applied to the membrane (Fig. 3C). Using Eq. S13b, we calculated the volume of the obtained shape at F_dimple_ = 3.73 pN/*μ*m^2^. We found that the volume of the simulated RBC at F_dimple_ = 3.73 pN/*μ*m^2^ is about V = 76.78 *μ*m^3^ which in comparison with Fig. 3C is closer to the reported experimental value for the RBC volume by Evans and Fung [69]. From these results, we can conclude that there is a better agreement between the simulated shape and the parametric shape of an experimentally observed RBC when a localized force is applied at the RBC dimple compared to the case with a uniform force distribution (Fig. 3).

### 3.3 Non-uniform distribution of force density in the RBC dimple region versus the rim region minimizes the shape error

While localizing the force density at the dimple decreased the error and the volume mismatch in our simulated RBC shapes, NMIIA is known to be distributed throughout the RBC [45]. Therefore, we next asked if the shape error can be minimized by including a normal force at the rim region in addition to the applied force at the dimple region (Fig. 5A). This analysis will allow us to predict the RBC shape not only in terms of absolute values of forces in the dimple and rim regions but also as a function of force per unit volume ratio in these two regions. In our model, based on the given force density per unit area in the dimple (F_dimple_) and rim (F_rim_) regions, we define the ratio of forces per unit volume as

**Figure 5:**
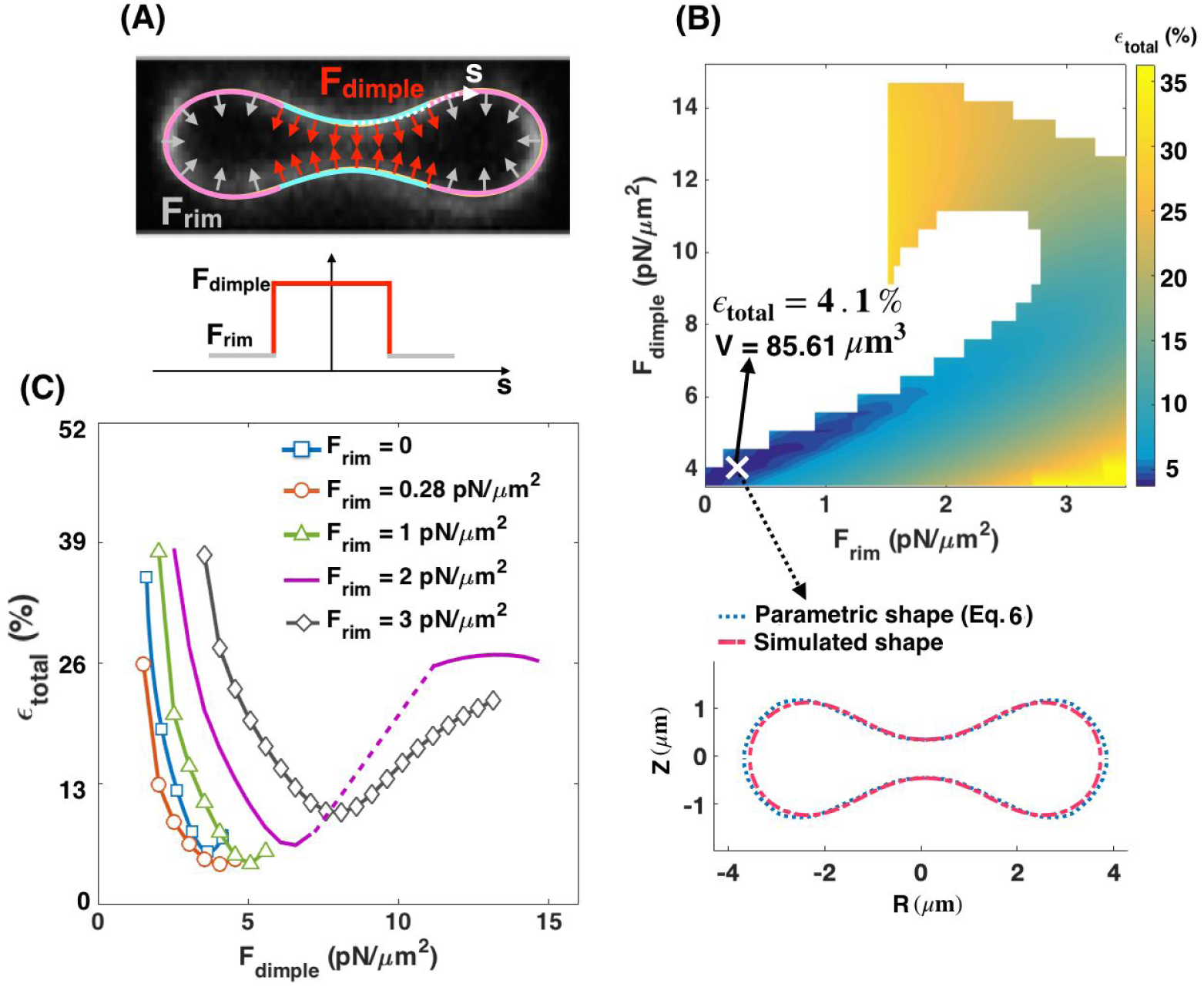
The applied force densities at the RBC dimple and rim regions regulate the shape error. (A, upper) Schematic of a biconcave RBC with a large force density (red arrows) at the dimple and a small force density (gray arrows) at the rim region. Schematic is overlaid on an RZ view of the center of an RBC from a confocal Z-stack of an RBC stained for the membrane marker glycophorin A. (A, lower) The applied force density along the membrane as a function of the arclength (Eq. S23). (B) Heat map shows the calculated shape error (Eq. 8) for a range of the force densities at the dimple (F_dimple_) and rim (F_rim_) regions. We stopped the simulations when the height at the dimple tends to zero (h_min →_ 0). The marked point **X** shows the case that has the lowest value of the error in the heat map at F_dimple_ = 4.05 pN/*μ*m^2^ and F_rim_ = 0.28 pN/*μ*m^2^ (ϵ_total_ ∼ 4.1%) with V = 85.61 *μ*m^3^. A comparison between the parametric shape of an RBC (dotted blue line) and the shape obtained from the simulation at point **X** (dashed red line) is shown in the lower panel. (C) The shape error as a function of force density at the dimple (F_dimple_) for five different values of the applied force density at the rim region. The dotted purple line shows a discontinuous transition in the shape error with increasing the dimple force density for F_rim_ = 2 pN/*μ*m^2^. Similar to Fig. 4B, independent of the value of F_rim_, the total error is a nonlinear function of the dimple force density (F_dimple_).

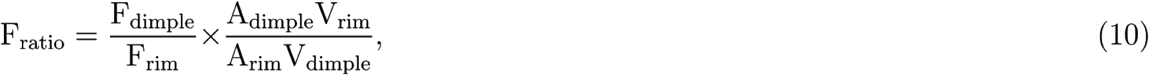

where A_dimple_ and A_rim_ are the area of the membrane surface in the dimple and rim regions, and V_dimple_ and V_rim_ are the volume occupied by the dimple and rim regions, respectively. For a given RBC shape, the area and the volume of the dimple and rim regions can be calculated by Eq. S13a and Eq. S13b, respectively.

We begin our analysis with the case where the pulling force in the dimple area is larger than the pulling force in the rim area (F_rim_ < F_dimple_). We implemented this distribution of force along the RBC membrane via a hyperbolic tangent function (Eq. S23) and performed the simulations over a range of forces at the dimple and the rim regions (F_dimple_ = 3.5 - 14 pN/*μ*m^2^ and F_rim_ = 0 - 3.5 pN/*μ*m^2^). The range of dimple force (F_dimple_) is chosen based on our previous results (Fig. 4) to have a close comparison with the parametric shape and obtain biconcave shapes from simulations with h_max_> h_min_ and h_min_> 0. The force along the rim (F_rim_) is set between F_rim_ = 0 and F_rim_= 3.5 pN/*μ*m^2^ based on the imposed condition of F_ratio_ <1 for all simulations.

The heat map in Fig. 5B represents the magnitude of the shape error for a given force density at the dimple and rim area. The simulations were stopped when the height at the RBC dimple approached zero, shown as white domains in the heat map (Fig. 5B). Based on these calculations, we found that the shape error has the lowest value (ϵ_total_ ∼ 4.1%) when F_dimple_ = 4.05 pN/*μ*m^2^ and F_rim_ = 0.28 pN/*μ*m^2^ (the point **X** on the heat map). For these specific force values, the parametric shape of the RBC (Eq. 6) and the shape obtained from the simulation at point **X** are very well-matched (Fig. 5B lower panel). Additionally, the volume of the simulated shape at point (**X**), is close (V = 85.62 *μ*m^3^) to the experimentally reported value by Evans and Fung [69].

To further understand the relationships between F_rim_ and F_dimple_ in governing the shape of the RBC, we plotted the shape error as a function of F_dimple_ for five different values of the force density at the rim section (Fig. 5C). We found that the shape error shows the same nonlinear dependence for different values of F_rim_. By increasing the value of F_dimple_, the shape error initially decreases by an order of magnitude and attains a relative minimum for each curve (Fig. 5C). Any further increase in the dimple force density results in a larger shape error (Fig. 5C), similar to Fig. 4B. As expected from Fig. 5B, the shape error has the lowest value on the red curve (F_rim_ = 0.28 pN/*μ*m^2^) when F_dimple_ = 4.05 pN/*μ*m^2^. Using Eq. 10, this set of dimple and rim forces in Fig. 5B is equivalent to F_ratio_∼ 14.27, which reflects the fact that to obtain the best match between the simulated RBC shape and the experimentally observed morphology, 14.27 times larger force per unit volume should be applied in the dimple region than the rim region.

Thus far, we have only considered the cases in which NMIIA motors were able to exert small pulling forces in the rim region. However, two other force configurations are possible; (i) NMIIA motors apply a larger force density in the rim region than the dimple area (F_rim_ > F_dimple_) (Fig. S2A), and (ii) NMIIA motors exert pushing forces in the rim region (Fig. S3A). We found that a large pulling force in the rim region (F_rim_ > F_dimple_) generates a shape resembling a peanut-shaped vesicle with a large shape error of ϵ_total_ >> 50% (Fig. S2). We also observed that applying a pushing force in the rim region (F_rim_ = 3.73 pN/*μ*m^2^) with no force in the dimple causes an error of ϵ_total_ ∼ 12.5% (Fig. S3B). Even adding a small pushing force in the rim region (F_rim_ = 0.53 pN/*μ*m^2^) with F_dimple_ = 3.73 pN/*μ*m^2^ increases the shape error to ϵ_total_ ∼ 9.7% (Fig. S3C). Our major prediction is that RBC biconcave shape depends on a heterogeneous distribution of NMIIA forces, which can be accomplished by more NMIIA motors density in the dimple compared to the rim.

### 3.4 RBC dimple region has a higher concentration of the NMIIA puncta as compared to the rim region

Our simulations suggest that NMIIA-mediated force densities are not uniformly distributed across the RBC membrane but instead are larger in the dimple region than the rim region (Fig. 5). Therefore, we hypothesized that the NMIIA distribution in RBCs is also non-uniform, with more NMIIA in the dimple region than the rim region. To test this hypothesis, we localized NMIIA motor domain puncta in three-dimensional reconstructions of AiryScan confocal Z-stacks [45,93]. This assay detects myosin bipolar filaments and other higher-order structures, as individual NMIIA molecules are too dim to detect in AiryScan images.

We divided each RBC into dimple and rim regions based on F-actin staining at the membrane (Fig. 6A) and quantified the number of NMIIA motor domain puncta in each region and the volumes of each region and the whole RBC using Volocity software. The dimple region accounted for about 7.4% of the total RBC volume (based on the F-actin staining, Fig 6B). This value agrees with calculations of dimple volume (∼7.1% of total volume) from our simulated shapes, in which we classify the dimple and rim regions based on the sign of the local mean curvature (Fig. 1B). The number of NMIIA puncta varies between RBCs, with 125 ± 47 puncta in the whole RBC, 113 ± 42 puncta in the rim, and 12 ± 9 puncta in the dimple (all values are mean ± SD). The dimple region contains about 9.1% of the total NMIIA motor domain puncta (Fig. 6C). In the dimple and rim regions as well as the whole RBC, the number of NMIIA puncta tends to increase with increasing the region or cell volume (Fig. S4).

**Figure 6:**
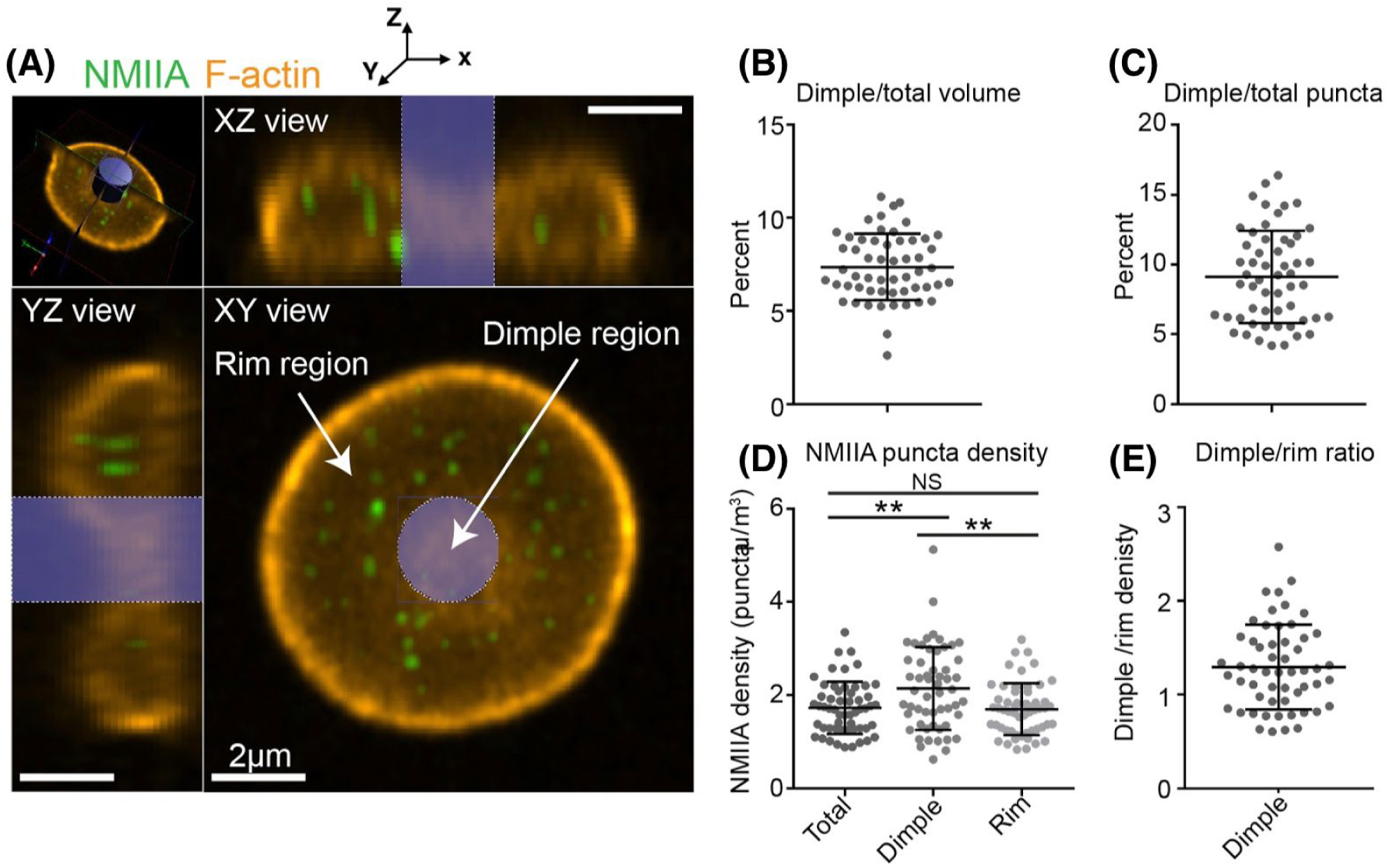
The RBC dimple has a higher average NMIIA puncta density than the RBC rim. (A) Optical section of a super-resolution Airyscan confocal Z-stack of human RBC immunostained with an antibody to the motor domain of NMIIA (green) and rhodamine-phalloidin for F-actin (orange). The top left image shows a perspective view of the optical section. Top right and bottom left images show YZ and XZ slices, respectively, of the RBC from planes perpendicular to this optical section. The bottom right image shows an XY view of the optical section. The blue cylinder represents the region identified as the dimple region. The rest of the RBC is identified as the rim region. Note, the myosin puncta near the RBC membrane are difficult to visualize in these merged images due to the bright F-actin staining. (B) The percent of total RBC volume occupied by the dimple region. Mean ± S.D. = 7.37 ± 1.79. (C) The percent of total NMIIA puncta in the dimple region. Mean ± S.D. = 9.11 ± 3.30. (D) The RBC dimple region has a ∼25% higher density of NMIIA puncta than whole RBCs (Total) (p = 0.0051) or the rim region (p = 0.0023) by Tukey’s multiple comparisons test. Mean ± S.D.: Total = 1.73 ± 0.562; Dimple = 2.15 ± 0.888; Rim = 1.70 ± 0.556. (E) Ratio of dimple and rim region NMIIA puncta densities for each RBC. Mean ± S.D. = 1.29 ± 0.452. (B-E) n = 55 RBCs from 3 individual donors.

The number of NMIIA puncta per unit volume (*μ*m^3^) in an RBC region is likely proportional to the number of NMIIA filaments that interact with membrane skeleton F-actin to exert force on the RBC membrane. The whole RBC and the rim region have similar NMIIA puncta densities (1.73 ± 0.562 *μ*m^3^ and 1.70 ± 0.556 *μ*m^3^, respectively), while the dimple region has a ∼25% higher density (2.15 ± 0.888 *μ*m^3^) (Fig. 6D). Thus, the dimple region has ∼1.29 times higher NMIIA puncta density compared to the rim region (Fig. 6E).

To determine whether differences in NMIIA densities relate to the extent of RBC biconcavity, we related NMIIA density to the minimum and maximum heights of XZ slices at the center of each RBC (Fig. S5). In both whole RBCs (Fig. S5A) and the dimple region (Fig. S5C), RBC biconcavity increased with increasing NMIIA density, while NMIIA density in the rim region was not related to biconcavity (Fig. S5B). These results agree with the results of our simulations, which predict that the maximum height of the rim (h_max_) is the least critical dimension to minimize the shape error (Fig. 3) and furthermore, that NMIIA exerts a larger force density at the RBC dimple (Fig. 5). Together, our simulations and experimental data suggest that this non-uniform force distribution is required to specify RBC biconcave disk shape.

### 3.5 Effective membrane tension regulates the required force densities ratio in the RBC dimple versus the rim region

We found that for the simulated RBC shapes, the shape error is minimized when the force per unit volume applied normally in the dimple region is about 14.27 times larger than the force per unit volume applied in the rim region (F_ratio_ = 14.27), in a tensionless membrane (Fig. 5). However, our experimental measurements reveal that in a healthy human RBC, the dimple region has only ∼25% higher density of NMIIA puncta than the rim region (Fig. 6). If we assume that the NMIIA density is proportional to the force generation capacity, then the induced force in the dimple region should be 1.25 times larger than the rim area. Therefore, we set out to reconcile this apparent discrepancy in the predicted F_ratio_ and measured the NMIIA density ratio. We found an interesting observation in the literature that the membrane tension in RBCs can vary from 10^−1^ pN/nm to 10^−4^ pN/nm [65,91,94]. Here, we interpret membrane tension to be the effective contribution of the membrane in-plane stresses and the membrane-cytoskeleton interactions [95]. We hypothesized that this in-plane tension of the RBC could play a critical role in relating the RBC shape to the NMIIA-generated force ratio in dimple and rim regions.

To investigate how this variation in membrane tension can modulate F_ratio_ and the shape error, we repeated the simulations as in Fig. 5 for three different effective membrane tensions: *(i)* low membrane tension (tension = 10^−4^ pN/nm) (Fig. 7A), *(ii)* intermediate membrane tension (tension = 10^−3^ pN/nm) (Fig. 7B), and *(iii)* high membrane tension (tension = 10^−2^ pN/nm) (Fig. 7C). The marker (**X**) in each heat map shows the point with minimum shape error for that set of simulations. To visualize the geometry of the simulated RBC at each point marked with an ‘**X’**, we plot the shapes that were obtained from simulations (solid yellow line) versus the reference experimental data (dotted blue line) [69] and also calculated the volume of the simulated geometry using Eq. S13b.

**Figure 7:**
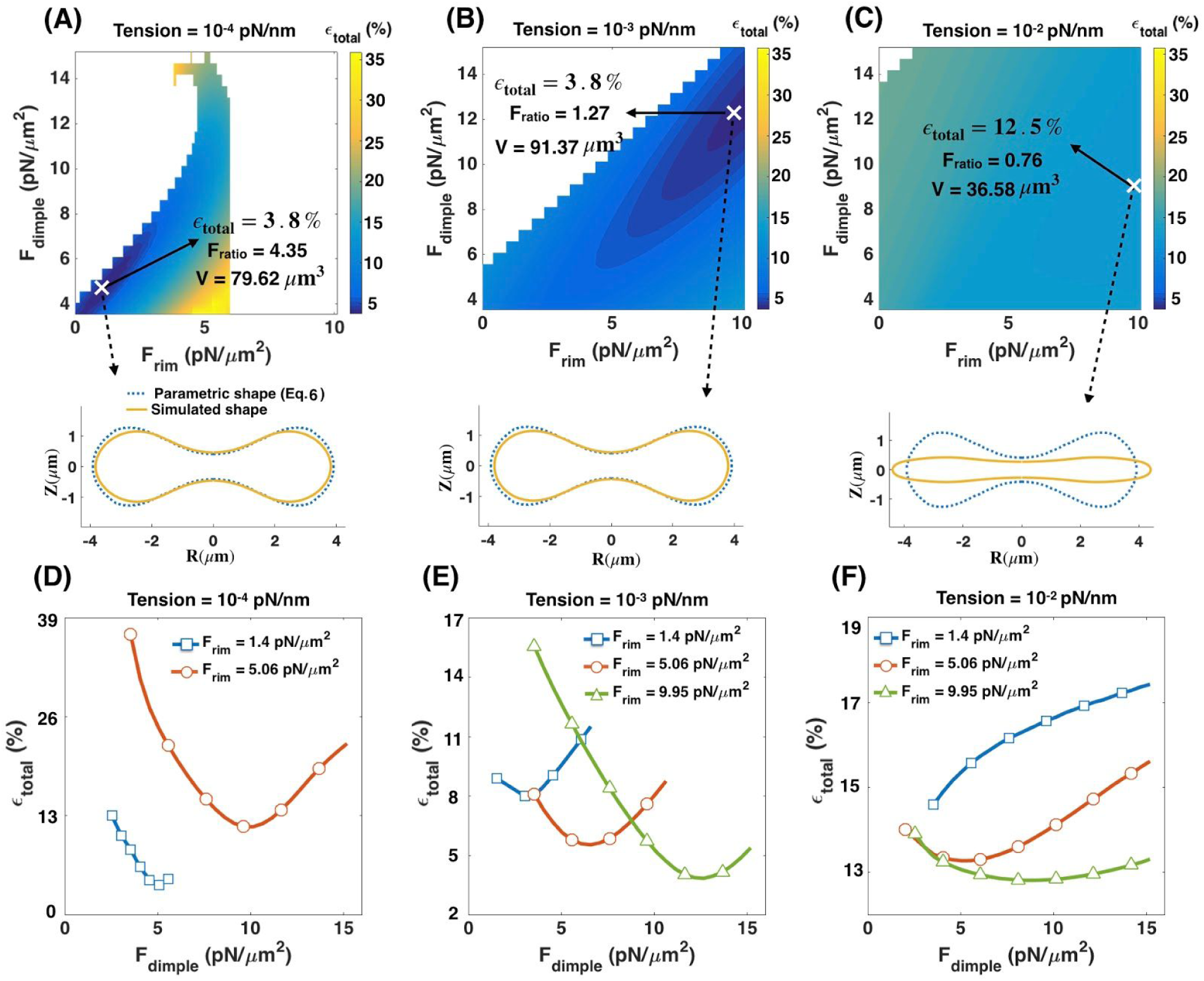
Effective membrane tension is a key parameter in regulating the RBC biconcave shape in addition to applied forces in the dimple and rim regions. (A-C) Heat maps show the total error in the shape of the simulated RBCs for (A) low tension (tension = 10^−4^ pN/nm), (B) intermediate tension (tension = 10^−3^ pN/nm), and (C) high tension (tension = 10^−2^ pN/nm). In each heat map, the point with the minimum error is marked with **X.** Also, for each marked point, the volume of the simulated RBC (V) is calculated using Eq. S13b, and the shape (solid yellow line) is shown in comparison with the reference parametric shape (dotted blue line). At intermediate tension, the shape error has the lowest value when F_ratio_ = 1.27 consistent with our experimental results in Fig. 6. (D-F) The calculated shape error (Eq. 8) as a function of the dimple force density (F_dimple_) for different values of the force density at the rim region and the membrane tension.

We observe that the shape error remains almost constant (ϵ_total_ ∼3.8%) with increasing the membrane tension from zero to low and intermediate values (Fig. 7A, B). However, varying the membrane tension alters the force ratio that gives the minimum shape error as well as the volume of the simulated geometry. For example, at low tension, the minimum shape error occurs at F_ratio_ = 4.35 where V = 79.62 *μ*m^3^ and at intermediate tension the shape error is minimum when F_ratio_ = 1.27 with V = 91.37 *μ*m^3^ (Fig. 7A, B), close to the volume experimentally reported by Evans and Fung [69]. In the case of high membrane tension, we found that the simulated shape deviates significantly from the biconcave disk and becomes closer to a pancake with a small volume (V = 36.58 *μ*m^3^) and the error goes up noticeably to about 12 percent (ϵ_total_ ∼ 12.5%) (Fig. 7C).

Additionally, we found that for low and intermediate tensions independent of the value of F_rim_, the shape error has the same non-linear relationship with increasing F_dimple_ as previously observed for the tensionless membrane (Fig. 7D, E). At low tension, the minimum shape error occurs when F_dimple_ = 5.06 pN/*μ*m^2^ and F_rim_ = 1.4 pN/*μ*m^2^ (blue square line) (Fig. 7D). At intermediate tension, a combination of F_dimple_= 12.66 pN/*μ*m^2^ and F_rim_ = 9.95 pN/*μ*m^2^ gives the minimum shape error (green triangle line) (Fig. 7E). However, for high tension, because of the stiffness of the membrane, we observe not only a deviation from the biconcave shape but also a deviation from the nonlinear error - dimple force relationships (Fig. 7F).

Based on these results we conclude that in addition to a non-uniform force distribution along the RBC membrane, a non-zero intermediate tension is required to obtain a close match between the shape and the volume of the simulated RBC and the experimental data. Furthermore, the intermediate value of tension (tension = 10^−3^ pN/nm) gives an excellent quantitative match for the predicted value of F_ratio_ (Fig. 7B) and the experimentally observed NMIIA density ratio (Fig. 6).

### 3.6 The angle of applied forces in the RBC dimple and rim regions controls the shape error

Until now, we have assumed the net effects of NMIIA motor proteins act as local forces applied normally to the membrane surface. However, there is evidence that these molecules also exert forces tangential to the membrane [96]. To examine how the orientation of the induced forces by NMIIA can affect the morphology of the RBC, we repeated the simulation in Fig. 7 for different membrane tension values assuming that the applied forces make an angle *ϕ* with the tangent vector **a**_**s**_ (Fig. 8A). Because the exact orientation of the applied forces by NMII molecules is currently unknown, we varied angle *ϕ* from *ϕ* = 90° (normal to the membrane) to *ϕ* = 0 (tangential to the membrane) and for each case found the combination of the force densities that gives the minimum shape error (Figs. S6-S9).

**Figure 8:**
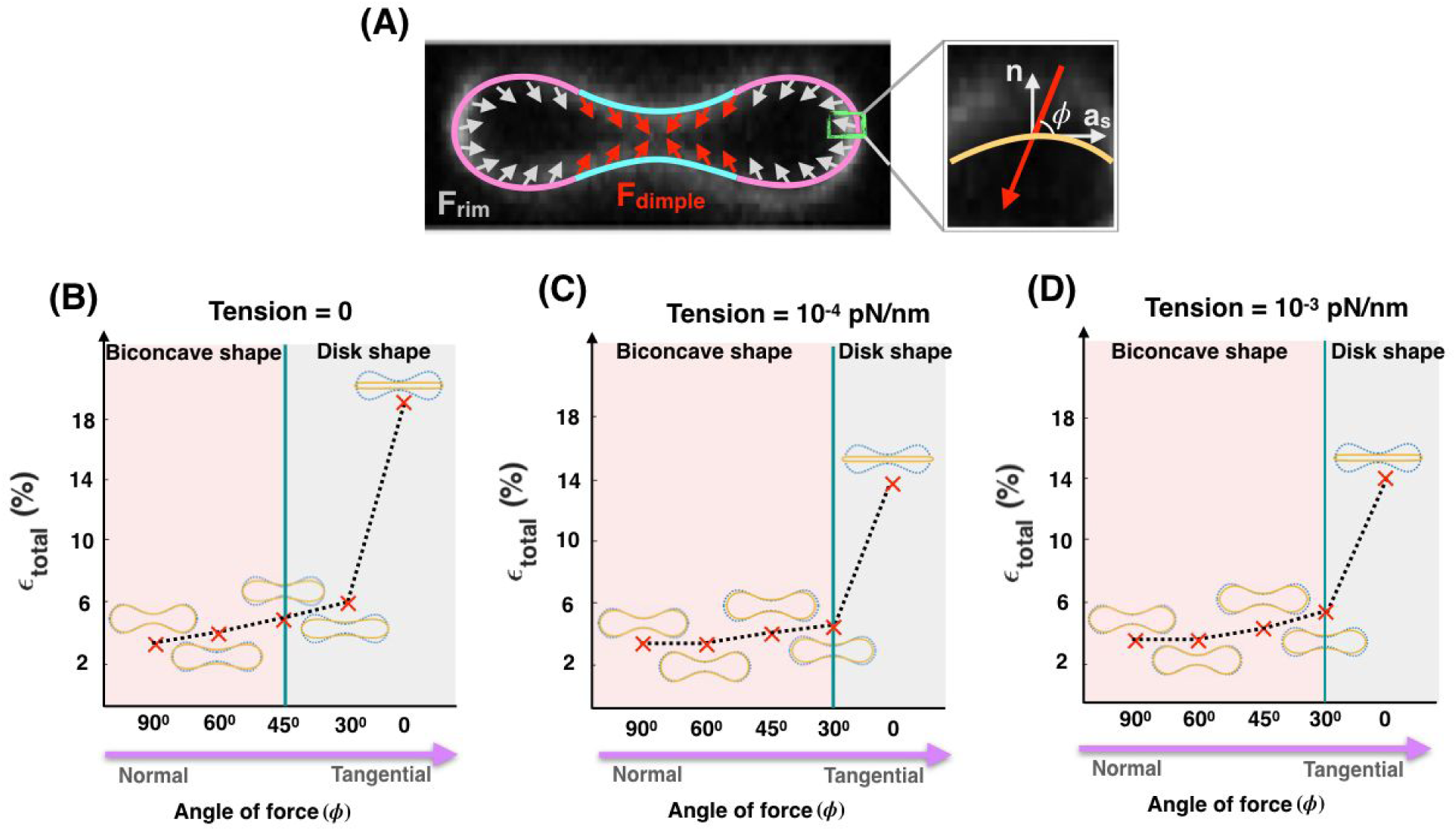
Effective membrane tension and the angle of applied forces in the RBC dimple and rim regions work together to maintain the biconcave shape of an RBC. (A) Schematic of a biconcave RBC with a non-uniform distribution of force density across the dimple and rim regions. In both regions, the forces per unit area are applied with angle *ϕ* with respect to the tangent vector (**a**_**s**_). (B-D) The shape error and the RBC shapes obtained from simulation for different angles of the applied forces (*ϕ*) for (B) tensionless membrane, (C) low tension (tension = 10^−4^ pN/nm), and (D) intermediate tension (tension = 10^−3^ pN/nm). For all values of the membrane tension, as the angle of forces deviates from normal (*ϕ* = 90°) to tangential orientation (*ϕ* = 0), the simulated shapes flatten and the shape error increases.

In Figs 8B-D, we plotted the minimum shape error as a function of angle *ϕ* for three different values of the membrane tension; (B) tensionless membrane, (C) low membrane tension (tension = 10^−4^ pN/nm), and (D) intermediate tension (tension = 10^−3^ pN/nm). We observed that in all three cases, with varying the angle *ϕ* from normal to tangential orientation, the simulated shapes deviate from the biconcave disks to the pancake shapes with an almost three times larger shape error. Based on our results, we found that for tensionless membranes, the transition from the biconcave shapes with ϵ_total_ < 5% (pink area) to the pancake shapes where ϵ_total_ >> 5% (gray area) occurs for angles smaller than 45° (*ϕ* < 45°) (Fig. 8B). This transition to pancake shapes shifted toward the smaller angles (*ϕ* < 30°) for low and intermediate membrane tension (Fig. 8C, 8D). Finally, in the case of high membrane tension, as expected from Fig. 7C, independent of the angle of force *ϕ*, the simulated shapes have a pancake-shaped morphology with very large shape errors (see Fig. S9). These findings suggest that membrane tension and the orientation of the applied forces can be tuned to actively maintain the biconcave morphology of an RBC.

## 4 Discussion

The biconcave disk shape of mammalian RBCs provides a maximum surface-area-to-volume ratio, which enables efficient gas and ion exchange and increases RBC deformability and resiliency [97]. This shape has been studied extensively from a mechanical standpoint to identify stress-strain relationships in cell membranes. Most studies modeling RBC shapes have been based on the work of Canham and Helfrich [53,59] and have reinforced the idea that mechanical force balance on the membrane by itself can provide insights into the unique shape of the RBC. The studies by Canham and Helfrich and other researchers suggested that the minimization of the membrane bending energy and the asymmetry between the inner and outer membrane leaflets generate the RBC biconcavity [34,35]. For example, Markin showed how the induced nano-scale curvature field due to the lateral distribution of membrane components can prescribe the biconcave shape of RBCs [37]. Here, we focused on micron-scale differences in lateral distribution of myosin-mediated forces as another degree of freedom and demonstrated how they are important for maintaining the RBC biconcave shape.

In RBCs, the skeleton underneath the plasma membrane is an elastic network of spectrin linked to short actin filament nodes that are attached to the membrane by anchoring proteins [30,31]. Studies of human and mouse congenital hemolytic anemias have also established a role for the RBC membrane skeleton in maintaining RBC biconcave shape in circulation [23,24,30]. Many different studies have demonstrated the importance of membrane/skeleton interactions in the formation of unusual RBC shapes as well as RBC deformability in shear flow [98–101]. In these studies, the RBC lipid bilayer and the membrane skeleton were mainly modeled as two distinct elastic components connected to each other via bond proteins [40–42]. However, no theoretical models for RBC biconcave shape have considered an active role for mechanochemical forces due to myosin motor proteins interacting with membrane skeleton F-actin in regulating RBC morphologies [40,102].

A recent study by Smith et al. [45] highlighted a critical role for NMIIA interactions with F-actin in the membrane skeleton in controlling RBC membrane tension and curvature. This discovery of a new experimental phenomenon calls for explanation and new/modified models that explicitly incorporate the effects of the molecular motors into the models based on pure membrane mechanics. Ultimately, RBCs can be used as a simple model system to explore the general role of NMII-generated forces in regulating plasma membrane curvature since RBC are the only cell type where F-actin is exclusively in the membrane skeleton [30]. The ubiquity of the membrane skeleton at the plasma membranes of all metazoan cells, where F-actin is also present in a transcellular cytoskeleton, further emphasizes the utility of the RBC paradigm.

In this study, we revisited the classical Helfrich-Canham energy model for the RBC membrane to include non-uniform forces along the membrane due to NMIIA-actin interactions. Undoubtedly, adding an additional degree of freedom to the energy allows us to attain a better match between the simulated and the experimentally observed RBC shapes compared to previous studies. Further, based on our results, we identify two conditions that need to be satisfied to produce the best fit with the experimental shapes of RBCs. First, the density of the NMIIA-generated force must be non-uniform along the RBC membrane to produce the best fit with the shapes measured experimentally. By conducting a parameter sweep of the force density configurations, we found that the non-uniform force distribution must be such that F_dimple_ is larger than F_rim_ (Figs. 4, 5). Experimental measurements of NMIIA density in the dimple and rim regions of RBCs using immunofluorescence showed that indeed NMIIA density is higher in the dimple than in the rim (Fig. 6) by about 25%. Our combined computational and experimental results highlight that a micron-scale, non-uniform force distribution of NMIIA plays a fundamental role in maintaining the biconcave shape of RBCs. We emphasize that this non-uniform density of forces is at the length scale of microns rather than at the length scale of fluctuations of the RBC membrane [103].

Second, we found that the effective membrane tension and the orientation of the applied forces are important physical parameters in modulating the RBC morphology and the required NMIIA-mediated force density ratio in the RBC dimple versus the rim region (F_ratio_) (Figs. 7, 8). As compared to tensionless or low-tension membranes, the intermediate tension values F_ratio_ for minimum shape error (∼1.27) are a better match with the experimentally reported NMIIA density ratio at the dimple versus the rim. Furthermore, we found that deviation of the applied forces from normal to tangential orientations results in pancake-shaped morphologies with very large shape errors compared to the actual biconcave shape of RBCs (Fig. 8).

Therefore, we predict that in mature, healthy biconcave RBCs, NMIIA motor domains exert force on the membrane with angle *ϕ* > 30° under intermediate membrane tension (∼10^−2^ pN/nm). The exact value of membrane tension and the angle of forces in an intact RBC are hard to measure because of the contributions from both the membrane and the underlying skeleton [104,105]. In the literature, a wide range of values are reported for the membrane tension from 10^−1^ pN/nm to 10^−4^ pN/nm [65,91,94]. This range can be attributed to dynamic lipid rearrangements [106], membrane-skeleton interactions [107], and based on our work here, rearrangement of force-generating NMIIA molecules [45]. The angle of applied forces by NMIIA at the RBC membrane, is still a matter of debate because nanoscale 3D images of F-actin and NMIIA motors using cryoelectron tomograms would be required to explore the relative configurations of myosin motors, F-actin and the RBC membrane surface[105]. This will require the development of novel sample preparation approaches for RBCs and is a subject for future study. Our theoretical analyses, supported by experimental measurements, implicitly suggest that for a biconcave RBC, the effective membrane tension should be on the order of 10^−2^ pN/nm and the NMIIA motors should apply forces with angles *ϕ* > 30° with respect to the membrane surface.

Our conclusions of non-uniform force density and tension regulation can be used to obtain insight into the effective activity of NMIIA motor domains at any given time. Assuming that a single NMIIA motor domain produces an average force of ∼ 2 pN [108,109], the calculated force densities in Fig. 7B correspond to 90 and 815 myosin motor domains in the dimple and rim regions, respectively. This means that the force generated by a total of ∼850 active NMIIA motor domains, distributed between the dimple and the rim as we predicted, is sufficient to sustain the biconcave disk shape of an RBC. Previous studies estimated that each mature human RBC contains ∼ 6,000 NMIIA molecules, ∼12,000 motor domains [43,44] and at any given time, roughly 40-50% of these molecules are bound to the membrane skeleton [45]. Our calculations suggest that approximately 15% of these bound NMIIA molecules are active and exerting forces distributed unevenly along the membrane. It is also possible that the amount of force generated by a single NMIIA motor domain varies due to the stiffness of the membrane skeleton network, the processivity (the duration over which the motor stays attached to actin), and the cross-linking activity of NMIIA myosin filaments [108,109]. Therefore, further research will be required to determine the quantitative relationship between the copy number of NMIIA molecules and their activity, that together determine the overall magnitude of the force exerted on the RBC membrane.

The idea of the asymmetrical distribution of the membrane skeleton and its components in the dimple and rim areas of RBCs was initially introduced by Hoffman, although no direct evidence for this was obtained [110,111]. Recently, Svetina et al. modeled RBC volume regulation according to the permeability of the Piezo1 channel. Based on their simulation results, they found that Piezo1 channels are expected to be distributed non-uniformly in a biconcave RBC, tending to localize in the dimple region[112]. They speculated that the simulated localization of Piezo1 channels in the dimple region is controlled by the membrane curvature. The RBC membrane curvature may also influence the localization of NMIIA motor proteins, as has been observed in other cell types [15]. Alternatively, a shear-induced Ca^2+^ influx through localized Piezo1 channels could locally activate NMIIA through phosphorylation of the regulatory light chain, leading to enhanced NMIIA binding to F-actin and enhanced local contractility at the dimple, activating Piezo1 and Ca^2+^ influx in a feed-forward loop. We believe our findings here are motivation for future studies to develop quantitative relationships between the myosin-mediated forces, Ca^2+^ influxes, and the membrane curvature of the cell surface.

We acknowledge that despite the conclusions from our studies, there are some limitations and simplifying assumptions that will need to be revisited for future studies. First, we limited our model to axisymmetric shapes, while RBCs often adopt non-axisymmetric shapes[113]. Future studies will involve simulations without any assumptions of symmetry. Experimental tests probing whether NMIIA activity is non-uniform along the RBC membrane will also give insight into NMIIA density distribution versus activity distribution along the membrane. Second, we assumed that the contributions from thermal fluctuations and the deformation of the membrane skeleton are negligible compared to the bending energy[55,114]. However, for a more general quantitative model, these effects should be considered [103]. Finally, future efforts focusing on the shape transformations of RBCs from discocytes to echinocytes or stomatocytes will be important to connect RBC morphology to physiological function and molecular mechanisms.

## Materials and methods

a. **Immunofluorescence staining of RBCs.** Human peripheral whole blood was collected from healthy human donors into EDTA tubes (BD Diagnostics). 20μl of whole blood was added to 1 ml of 4% paraformaldehyde (PFA, Electron Microscopy Sciences) in Dulbecco’s PBS (DPBS – Gibco), mixed, and incubated at room temperature overnight.
  i. **NMIIA immunostaining and rhodamine phalloidin staining.** Fixed RBCs were washed three times in DPBS by centrifuging for 5 minutes at 1000 x *g*, permeabilized in DPBS + 0.3% TX-100 for 10 minutes, and then blocked in 4% BSA, 1% normal goat serum in DPBS (Blocking Buffer, BB) at 4°C for at least 4 days or up to 1 week before immunostaining. Permeabilized and blocked RBCs were then incubated with rabbit anti-NMIIA motor domain antibody (Abcam ab75590) diluted in BB (1:1000) for 2-3 hours at room temperature, washed two times in BB as above, and then incubated in Alexa-488-conjugated goat anti-rabbit secondary antibody (Life Technologies A11008, diluted 1:1000) mixed with rhodamine-phalloidin (Life Technologies R415, at a final concentration of 130nM) in BB for 1-2 hr at room temperature, followed by washing three times in BB as above. Stained cells were cytospun onto slides and mounted with ProLong™ Gold mounting medium (Invitrogen) and coverslipped prior to imaging.
  ii. **Glycophorin A (GPA) immunostaining.** Fixed RBCs were washed three times in DPBS by centrifuging for 5 minutes at 1000 × *g*, blocked for 1 hour in BB, and stained with FITC-conjugated mouse anti-GPA antibody (BD Pharmingen 559943) for 1 hour at room temperature. GPA-stained RBCs were washed twice in DPBS by centrifugation as above, then cytospun onto glass slides and mounted with Prolong™ Gold and coverslipped prior to imaging.
b. **Fluorescence microscopy.**
  i. **RBCs immunostained for NMIIA and rhodamine phalloidin for F-actin.** RBCs were imaged using a Zeiss LSM 880 Airyscan laser scanning confocal microscope with a 63× 1.46 NA oil Plan Apo objective. Z-stacks were acquired at a digital zoom of 1.8 and a Z-step size of 0.168 μm. The distance between Z-steps was set to 0.10 μm in images used for NMIIA puncta analysis in Volocity (Quorum Technologies).
  ii. **RBCs immunostained for NMIIA and rhodamine phalloidin for F-actin.** RBCs were imaged using a Zeiss LSM 880 Airyscan laser scanning confocal microscope with a 63× 1.46 NA oil Plan Apo objective. Z-stacks were acquired at a digital zoom of 1.8 and a Z-step size of 0.168 μm.
  iii. **RBCs immunostained for GPA.** RBCs were imaged using a Zeiss LSM 780 laser scanning confocal microscope with a 100× 1.4 NA oil Plan Apo objective. Z-stacks were acquired at a digital zoom of 1.0 and a Z-step size of 0.25 μm. The distance between Z-steps was set to 0.18 μm in images used for presentation.
c. **Image analysis.** Numbers of NMIIA puncta in whole RBCs, in the dimples, and in the rims were counted automatically from Airyscan confocal stacks in Volocity (Quorum Technologies) using the “Find Spots” function in the “Measurements” module. The volumes of whole RBCs, the dimples, and the rims were measured from the rhodamine phalloidin (F-actin) fluorescence in Volocity using the “Find Objects” function, with gaps in staining filled using the “Close” function. RBC height measurements were acquired manually from XZ views of the center of each RBC in Volocity using the line function to measure the distance between the edges of fluorescent F-actin staining signal at the widest and narrowest regions of each RBC.
d. **Statistical analysis.** Data are presented in dot plots as mean ± standard deviation (SD), or in scatter plots showing the best-fit line from the linear regression. Differences between the variances of the two samples were detected using F-tests. Differences between means were detected using unpaired t-tests with Welch’s correction. When more than one comparison was made, differences between means were detected using one-way ANOVA followed by Tukey’s multiple comparisons test. Statistical significance was defined as p < 0.05. Statistical analysis was performed using GraphPad Prism 7.03 software.

## Acknowledgment

This work was supported by ONR N00014-17-1-2628 to P.R. V.M.F was supported by NIH grant HL083464. H.A. was supported by a fellowship from the Visible Molecular Cell Consortium (VMCC), a program between UCSD and the Scripps Research Institute. A.S.S. was supported by a fellowship from the NIH/National Center for Advancing Translational Sciences Clinical and Translational Science Award TL1 TR00113 to the Scripps Translational Science Institute.

## Author contributions

H.A, A.S.S, R.B.N, V.M.F and P.R. conceived the research, H.A. and P.R. conducted the modeling simulations. A.S.S, R.B.N, and V.M.F analyzed the experimental data, H.A, A.S.S, R.B.N, V.M.F and P.R. wrote the paper. All authors reviewed the manuscript and agreed on the contents of the paper.

