## Supplemental material for "Non-uniform distribution of myosin-mediated forces governs red blood cell membrane curvature through tension modulation" (from another version of this preprint: version 1, 2019-06-12)

**3 Supplementary figures** **8**

### 1 Model description

#### 1.1 Assumptions

- We consider that the curvature of the membrane is much larger than the thickness of the bilayer [1]. This allows us to treat the lipid bilayer as a thin elastic shell and model the bending energy of the membrane by the Helfrich–Canham energy, which depends only on the local curvatures of the surface [2, 3].
- Due to the high stretching modulus of lipid bilayers, we assume that the membrane is locally incompressible/inextensible [4]. We use a Lagrange multiplier to implement this constraint [5–7].
- We assume that the applied force by the NMIIA motor proteins is local and it is primarily normal to the membrane (see Fig. 1A) [8, 9]. The tangential contributions from the NMIIA-mediated forces are assumed to be negligible. This assumption can be justified based on the preferably orthogonal orientation of the short actin filaments to the erythrocyte membrane in resting conditions [10, 11].
- We assume that the RBC is at mechanical equilibrium at all times, allowing us to neglect inertia [12–14]. This assumption is consistent with the experimentally observed shapes for the resting RBCs in both *vivo* and *vitro* [8, 15].
- We assume that the total surface area of the RBC membrane is constant and is 135 $\mu\text{m}^2$ [16, 17]. All our simulations are conducted using this constant area.
- For simplicity in the numerical simulations, we assume that the RBC is rotationally symmetric and also has a reflection symmetry with respect to the Z = 0 plane (see Fig. 1C) [3, 16, 18, 19]. This assumption reduces the computational cost of the simulation into just one curve as shown by the red dotted line in Fig. 1C.

#### 1.2 Equations of motion

In this section, we present a concise derivation of the governing mathematical shape equations for the RBC membrane at mechanical equilibrium. The complete derivation with details are given in [5, 20–22].
At equilibrium, the strain energy of the whole system is written as the integration of local energy density ($W$) over the total membrane surface area $\omega$ [23–25]

$$E = \int_{\omega} W(H, K; \theta^{\alpha}) da, \tag{S1}$$

where $E$ is total strain energy, $H$ is the mean curvature of the surface, $K$ is the Gaussian curvature, and $\theta^{\alpha}$ denotes the surface coordinate where $\alpha \in \{1, 2\}$.

To impose the incompressibility constrain, we can rewrite the energy equation Eq. S1 using a Lagrange multiplier

$$E = \int_{\omega} [JW(H, K; \theta^{\alpha}) + \lambda(\theta^{\alpha})(J - 1)] dA, \tag{S2}$$

where $\lambda$ is a Lagrange multiplier and $J = \sqrt{a/A}$ is the local areal stretch due to mapping from a reference frame to the actual surface. Here the physical interpretation for this parameter is membrane tension

Using the variational approach to minimize the total strain energy of a homogeneous membrane (Eq. S2) gives us the so-called "shape equation"

$$p + F = \Delta \frac{1}{2} W_H + W_H(2H^2 - K) + 2H(KW_K - W) - 2\lambda H, \tag{S3}$$

where $\Delta(\cdot) = (\cdot)_{;\alpha\beta}a^{\alpha\beta}$ is the surface Laplacian where $(\cdot)_{;\alpha}$ denotes the covariant derivative and $a^{\alpha\beta}$ is the dual metric, $p$ is the pressure difference across the membrane, and $F$ is the force density (force per unit area) representing the normally applied force density to the membrane surface by the NMIIA motor proteins.

##### 1.3 Equations of motion in axisymmetric coordinates

We assumed the RBC has a rotationally symmetric shape and define the surface of revolution (Fig. 1C) by

$$\mathbf{r}(s,\theta) = R(s)\mathbf{e}_r(\theta) + Z(s)\mathbf{k}, \tag{S4}$$

where s is the arclength along the curve, R(s) is the radius from the axis of rotation, Z(s) is the height from the base plane, and $(\mathbf{e}_r, \mathbf{e}_\theta, \mathbf{k})$ form the basis coordinate. Defining $\psi$ as the angle made by the tangent with respect to the vertical gives

$$R'(s) = \cos(\psi), \quad Z'(s) = \sin(\psi), \tag{S5}$$

which satisfies the identity $(R')^2 + (Z')^2 = 1$, where $()'$ is the partial derivative with respect to the arclength. Using Eq. S5, we can define the normal ($\mathbf{n}$) and tangent ($\mathbf{a}_s$) vectors to the surface as

$$\mathbf{n} = -\sin\psi\mathbf{e}_r(\theta) + \cos\psi\mathbf{k}, \quad \mathbf{a}_s = \cos\psi\mathbf{e}_r(\theta) + \sin\psi\mathbf{k}. \tag{S6}$$

This parameterization allows us to write the tangential ($\kappa_\nu$) and transverse ($\kappa_\tau$) curvatures as

$$\kappa_\nu = \psi', \quad \kappa_\tau = \frac{\sin\psi}{R}. \tag{S7}$$

The mean curvature ($H$) and Gaussian curvature ($K$) are obtained by summation and multiplication of the tangential and transverse curvatures

$$H = \frac{1}{2}(\kappa_\nu + \kappa_\tau) = \frac{1}{2}(\psi' + \frac{\sin\psi}{R}), \quad K = \kappa_\tau\kappa_\nu = \frac{\psi'\sin\psi}{R}. \tag{S8}$$

Finally, we define $M = \frac{1}{2\kappa}R(W_H)'$ such that for a uniform bending rigidity ($\kappa$), we obtain a system of first-order differential equations with five unknowns $R$, $Z$, $\psi$, $H$, and $M$ [20, 26, 27],

$$\begin{gathered} R' = \cos\psi, \quad Z' = \sin\psi, \quad R\psi' = 2RH - \sin\psi, \quad RH' = M, \\ \text{and} \quad \frac{M'}{R} = \frac{p}{\kappa} + \frac{F}{\kappa} + \frac{2H\lambda}{\kappa} - 2H\left(H - \frac{\sin\psi}{R}\right)^2. \end{gathered} \tag{S9}$$

In order to solve the system of equations in Eq. S9, we need to provide five boundary conditions. We consider an axisymmetric RBC with reflection symmetry with respect to the Z = 0 plane (see Fig. 1C). These assumptions can be applied as the following boundary conditions,

$$\begin{gathered} R(0^+) = 0, \quad \psi(0^+) = 0, \\ Z(\mathrm{s_{max}}) = 0, \quad \psi(\mathrm{s_{max}}) = \frac{-\pi}{2}, \quad M(\mathrm{s_{max}}) = 0, \end{gathered} \tag{S10}$$

where $\mathrm{s_{max}}$ is the maximum length of the computational domain. One advantage of an asymmetric coordinate system is that the manifold area (A) and the occupied volume (V) can be expressed in term of arclength,

$$A(s) = 2\pi\int_0^s R(\eta)d\eta, \tag{S11a}$$

$$V(s) = 2\pi\int_0^s R(\eta)Z(\eta)d\eta, \tag{S11b}$$

which allows us to conserve the total area of the RBC by changing the maximum length of the computational domain ($s_{max}$).

##### 1.4 Nondimensionalization

In order to perform the numerical computations, we nondimensionalized the system of equations (Eq. S9) by using two positive constants, the radius of the RBC ($R_0$) and the lipid bilayer bending rigidity ($\kappa_0$). This allows us to define the dimensionless variables as

$$
\begin{gathered}
t = \frac{s}{R_0}, \quad r = \frac{R}{R_0}, \quad z = \frac{Z}{R_0}, \quad h = HR_0 \quad m = MR_0 \\
\tilde{\lambda} = \frac{\lambda R_0^2}{\kappa_0}, \quad \tilde{p} = \frac{pR_0^3}{\kappa_0} \quad f = \frac{FR_0^3}{\kappa_0}, \quad \tilde{\kappa} = \frac{\kappa}{\kappa_0}.
\end{gathered}
\tag{S12}
$$

Rewriting Eq. S9 in terms of the dimensionless variables, we get [27]

$$
\begin{gathered}
\dot{r} = \cos\psi, \quad \dot{z} = \sin\psi, \quad r\dot{\psi} = 2rh - \sin\psi, \quad x\dot{h} = m, \\
\frac{\dot{m}}{r} = \frac{\tilde{p}}{\tilde{\kappa}} + \frac{f}{\tilde{\kappa}} + \frac{2h\tilde{\lambda}}{\tilde{\kappa}} - 2h\left(h - \frac{\sin\psi}{r}\right)^2,
\end{gathered}
\tag{S13}
$$

where ( $\dot{}$ ) is the partial derivative with respect to $t$. With the defined dimensionless variables in Eq. S12, the boundary conditions simplified as

$$
\begin{gathered}
r(0^+) = 0, \quad \psi(0^+) = 0, \\
z(t_{max}) = 0, \quad \psi(t_{max}) = \frac{-\pi}{2}, \quad m(t_{max}) = 0.
\end{gathered}
\tag{S14}
$$

##### 1.5 Parametrization of RBC biconcave morphology and shape error estimation

There are several parametric models to describe the biconcave morphology of an RBC [28–31]. Initially, Funaki proposed the Cassini oval model for the RBC biconcave morphology given by [28]

$$
(R^2 + Z^2 + a^2) - 4a^2R^2 = c^4, \tag{S15}
$$

where $a$ and $c$ are constants with the condition that $a < c < \sqrt{2}a$. Yurkin modified the Cassini oval model to an implicit equation with four constants given by [29]

$$
R^4 + 2C_4R^2Z^2 + Z^4 + C_1R^2 + C_2Z^2 + C_3 = 0, \tag{S16}
$$

where $C_1$, $C_2$, $C_3$, and $C_4$ are constants that depended on the RBC dimension ($C_1 = -14.85$, $C_2 = 40.40$, $C_3 = -6.65$, and $C_4 = -0.30$). Borovoi et al. introduced a function that represents the biconcave shape of an RBC in the spherical coordinate as [30]

$$
R(\theta) = e\sin^q(\theta) + b, \tag{S17}
$$

where $e$, $b$, and $q$ are constants that were determined by fitting the function to the standard shape of an RBC ($e = 3\ \mu$m, $b = 0.75\ \mu$m, and $q = 5$). The most realistic model for the biconcave shape of an RBC was proposed by Evans and Fung based on direct experimental measurements [31]

$$Z(R) = \pm 0.5\sqrt{1-(\frac{2R}{L})}(D_1 + D_2(\frac{2R}{L})^2 - D_3(\frac{2R}{L})^4), \tag{S18}$$

where $D_1$, $D_2$, and $D_3$ are the constants that were calculated by fitting the function to the average dimension of an experimentally observed RBC and statistical analysis ($D_1 = 0.81\ \mu\text{m}$, $D_2 = 7.83\ \mu\text{m}$, and $D_3 = -4.39\ \mu\text{m}$). We plotted these different proposed parametric models together for the biconcave shape of an RBC for fixed $\text{h}_{\text{min}}$, $\text{h}_{\text{max}}$, and L (Fig. 2B). In this study, we used the given parametric function by Evans and Fung (Eq. S18) as the reference geometry for the RBC experimental shape.

Typically, there is a mismatch between each RBC shape obtained from our mechanical model (Eq. S13) and the RBC parametric shape (Eq. S18). Considering the three characteristic lengths ($\text{h}_{\text{min}}$, $\text{h}_{\text{max}}$, and L in Fig. 1C), we can define three errors that characterize the difference between each of these lengths in the simulated shapes and the reference experimental shape,

$$\begin{aligned} \epsilon_{\text{hmax}} &= |\text{h}_{\text{max,exp}} - \text{h}_{\text{max,sim}}| = |\Delta \text{h}_{\text{max}}| \\ \epsilon_{\text{hmin}} &= |\text{h}_{\text{min,exp}} - \text{h}_{\text{min,sim}}| = |\Delta \text{h}_{\text{min}}| \\ \epsilon_{\text{L}} &= |\text{L}_{\text{exp}} - \text{L}_{\text{sim}}| = |\Delta \text{L}|, \end{aligned} \tag{S19}$$

where $(.)_{\text{sim}}$ is the calculated length from the simulated shape and $(.)_{\text{exp}}$ is the measured length in the reference experimental shape. The total error ($\epsilon_{\text{total}}$) in the shape of the simulated RBC can be calculated by the root mean square (RMS) between each two mapped points of the simulated and parametric shapes

$$\epsilon_{\text{total}} = \sqrt{\frac{1}{N}\Big[\sum_{i=1}^{i=N}(Z_{i,\text{sim}} - Z_{i,\text{par}})^2 + (R_{i,\text{sim}} - R_{i,\text{par}})^2\Big]}, \tag{S20}$$

where $N$ is the total number of nodes across the RBC shapes, $i$ is the index node, $Z_{i,\text{sim}}$ and $Z_{i,\text{par}}$ are the height of the simulated and the RBC parametric (Eq. 3) shapes at index $i$, respectively. $R_{i,\text{sim}}$ is the radius of the simulated shape at index $i$, and $R_{i,\text{par}}$ is the radius of the RBC parametric shape (Eq. 3) at index $i$ (see Fig. 2C). To trace the simulated shape and the parametric shape (Eq. 3), we mapped the points with the maximum height, minimum height, and maximum radius. Then, we discretized the domain equally between the points and calculated the errors.

#### 1.6 Numerical implementation

We solved the system of first-order differential equations (Eq. S13) with boundary conditions Eq. S14 by the finite element software COMSOL MULTIPHYSICS ® 5.3a, using the “General Form PDE” module. Here, we summarize the steps and assumptions that we used for each simulation.

- All the simulations were performed for fixed total arclength. However, in each simulation, the maximum arclength ($\text{s}_{\text{max}}$) varies to conserve the total area of the RBC membrane.
- The computational domain ($t$) was discretized equally with mesh size = 0.001.
- To have a sharp but smooth transition in the distribution of the force ($f$), we used a hyperbolic tangent function given by

$$f = \frac{1}{2}[\tanh(g(t - s_{\text{dimple}}))], \tag{S21}$$

  where g is a constant (here we set g=20) and $s_{\text{dimple}}$ represents the length scale that the local force at the dimple is applied.

- The applied force in Figs. 3-5 and Fig. 7 was progressively increased such that each solution was used as an initial guess for the next step.

### 2 Tables

#### 2.1 Table of Notation

Table 1: Notation used in the model

| Notation | Description | Units |
|---|---|---|
| $p$ | Pressure difference across the membrane | $\mathrm{pN/nm^2}$ |
| $\theta^\alpha$ | The surface coordinate ($\alpha \in \{1,2\}$) | |
| $W$ | Local energy per unit area | pN/nm |
| $\mathbf{r}$ | Position vector | |
| $\mathbf{n}$ | Normal vector to the membrane surface | unit vector |
| $\mathbf{a_s}$ | Tangent vector to the membrane surface | unit vector |
| $\boldsymbol{\nu}$ | Tangent vector to the membrane surface in direction of increasing arclength | unit vector |
| $\boldsymbol{\tau}$ | Rightward normal in direction of revolution | unit vector |
| $\mathbf{a}_\xi$ | Basis vectors describing the tangent plane | |
| $\lambda$ | Membrane tension | pN/nm |
| $H$ | Mean curvature of the membrane | $1/\mathrm{nm}$ |
| $K$ | Gaussian curvature of the membrane | $1/\mathrm{nm}^2$ |
| $F$ | Normally applied force per unit area | $\mathrm{pN}/\mu\mathrm{m}^2$ |
| $\kappa_\nu$ | Tangential curvature | $1/\mathrm{nm}$ |
| $\kappa_\tau$ | Transverse curvature | $1/\mathrm{nm}$ |
| $\kappa$ | Bending modulus | $\mathrm{pN} \cdot \mathrm{nm}$ |
| $\kappa_G$ | Gaussian modulus | $\mathrm{pN} \cdot \mathrm{nm}$ |
| $s$ | Arclength | nm |
| $\theta$ | Azimuthal angle | |
| $\psi$ | Angle between $\mathbf{e}_r$ and $\mathbf{a}_s$ | |
| $R$ | Radial distance | nm |
| $Z$ | Elevation from base plane | nm |
| $\mathbf{e}_r(\theta)$ | Radial basis vector | unit vector |
| $\mathbf{e}_\theta$ | Azimuthal basis vector | unit vector |
| $\mathbf{k}$ | Altitudinal basis vector | unit vector |
| $A$ | Membrane area | $\mathrm{nm}^2$ |
| $s_{max}$ | Maximum arclength | nm |
| $\mathrm{h_{max}}$ | Maximum height at the rim | |
| $\mathrm{h_{min}}$ | Minimum height at the dimple | |
| L | The maximum cell diameter | |
| $\epsilon_{\mathrm{hmax}}$ | Error in the maximum height | |
| $\epsilon_{\mathrm{hmin}}$ | Error in the minimum height | |
| $\epsilon_{\mathrm{L}}$ | Error in the length | |
| $\epsilon_{\mathrm{total}}$ | Total error | |
| $M$ | Shape equation variable | $1/\mathrm{nm}$ |

Table 2: Notation used in the model

| Notation | Description | Units |
|---|---|---|
| $R_0$ | Radius of the RBC cell | nm |
| $\kappa_0$ | Bending rigidity of the lipid | pN |
| $\mathrm{F}_{\text{uniform}}$ | Uniformly applied force | $\text{pN}/\mu\text{m}^2$ |
| $\mathrm{F}_{\text{dimple}}$ | Local force at dimple region | $\text{pN}/\mu\text{m}^2$ |
| $\mathrm{F}_{\text{donut}}$ | Local force at donut region | $\text{pN}/\mu\text{m}^2$ |
| $\mathrm{A}_{\text{dimple}}$ | Membrane surface are in the dimple region | $\text{m}^2$ |
| $\mathrm{A}_{\text{donut}}$ | Membrane surface are in the donut region | $\text{m}^2$ |
| $\mathrm{V}_{\text{dimple}}$ | Occupied volume by the dimple region | $\text{m}^3$ |
| $\mathrm{V}_{\text{donut}}$ | Occupied volume by the donut region | $\text{m}^3$ |
| $r$ | Dimensionless radial distance | |
| $z$ | Dimensionless height | |
| $h$ | Dimensionless mean curvature | |
| $m$ | Dimensionless M | |
| $\tilde{\lambda}$ | Dimensionless surface tension | |
| $\tilde{p}$ | Dimensionless pressure | |
| $\tilde{f}$ | Dimensionless normal force | |
| $\tilde{\kappa}$ | Dimensionless bending modulus | |
| $t$ | Dimensionless arclength | |
| $\mathrm{F}_{\text{ratio}}$ | Ratio of the force at the dimple versus the donut region | |

#### 3 Supplementary figures

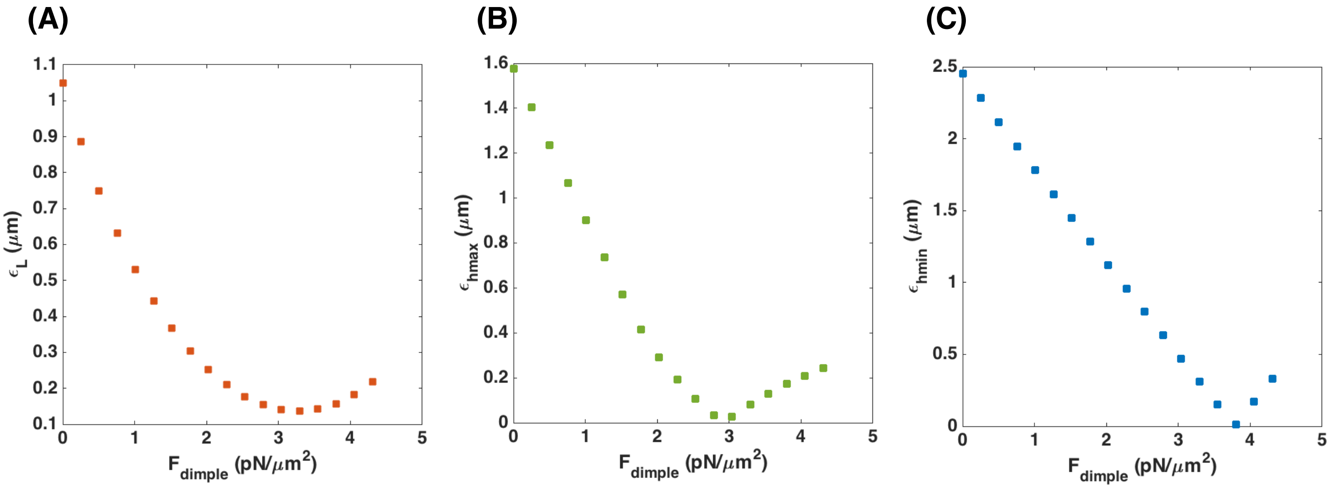

Figure S1: The error in the characteristic lengths (Eq. S19) is a nonlinear function of dimple force density ($F_{dimple}$). (A) Calculated error in the maximum length of the simulated RBC ($\epsilon_L$) as a function of $F_{dimple}$. (B) Calculated error in the maximum height of the rim of the simulated RBC ($\epsilon_{hmax}$) as a function of $F_{dimple}$. (C) Calculated error in the minimum height of the dimple of the simulated RBC ($\epsilon_{hmin}$) as a function of $F_{dimple}$. In all three graphs, with increasing $F_{dimple}$ from zero, initially the error decreases about an order of magnitude and attains a relative minimum. Any further increase in $F_{dimple}$ toward the large dimple force density ($F_{dimple} > 4\text{pN}/\mu\text{m}^2$) leads to a larger error in all characteristic lengths.

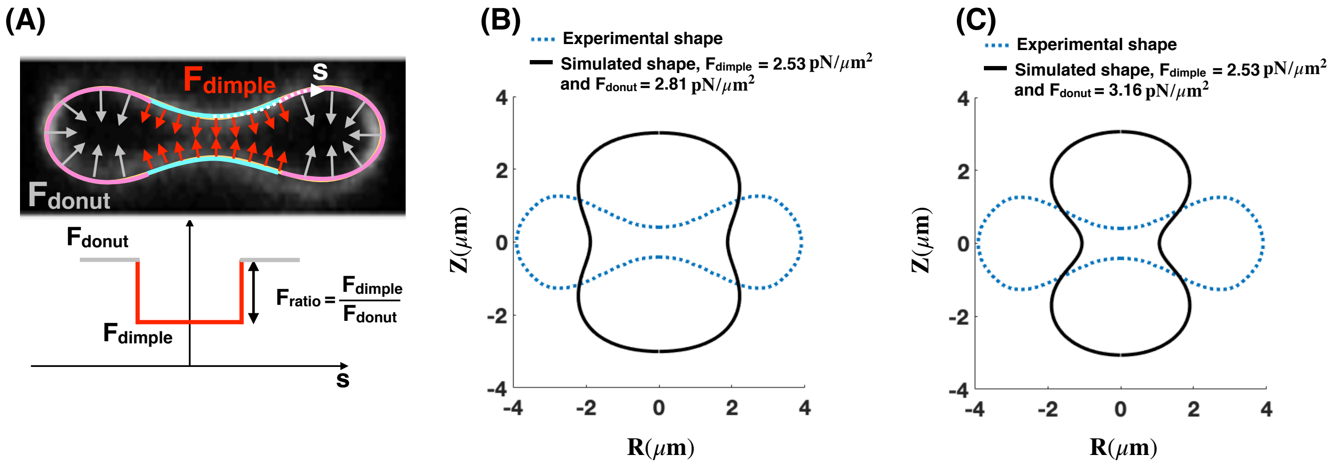

Figure S2: Applying a large pulling force at the donut region causes a very large shape error. (A, upper) Schematic of a biconcave RBC with a small force per unit area (red arrows) at the dimple and a large force per unit area (gray arrows) at the donut section. (A, lower) The applied force along the membrane as a function of the arclength (Eq. S21 ). (B) A comparison between the simulated shape of RBC with $F_{dimple}$ = 2.53 $\text{pN}/\mu\text{m}^2$ and $F_{donut}$ = 2.81 $\text{pN}/\mu\text{m}^2$ ($F_{ratio}$ = 0.9). Having a $F_{ratio} < 1$ condition deviates the simulated shape from a biconcave to a peanut-shape geometry. Therefore, the calculated shape error becomes significantly large ($\epsilon_{total} \sim 2.05\ \mu\text{m}$). (C) The reference experimental shape versus the shape that obtained from the simulation with $F_{dimple}$ = 2.53 $\text{pN}/\mu\text{m}^2$ and $F_{donut}$ = 3.16 $\text{pN}/\mu\text{m}^2$ ($F_{ratio}$ = 0.8). As the donut force density increases, we found a much larger shape error in the simulated geometry ($\epsilon_{total} \sim 3.56\ \mu\text{m}$).

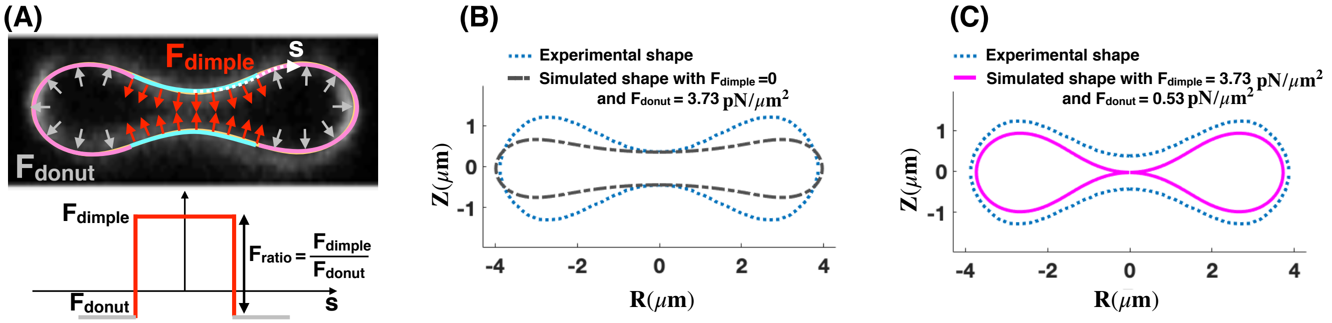

Figure S3: Applying a pushing force at the donut region increases the total shape error (Eq. S20). (A) (A, upper) Schematic of a biconcave RBC with an inward force per unit area (red arrows) at the dimple and an outward force per unit area (gray arrows) at the donut section. (A, lower) The applied force along the membrane as a function of the arclength (Eq. S21 ). The simulated shape of RBC with only a pulling force at the dimple region ( $F_{donut}$ = 3.73 $pN/\mu m^2$). Here, the total shape error is $\epsilon_{total} \sim 0.49\ \mu m$, which is larger than even the case with the uniform force per unit area (Fig. 3, center). (C) The simulated shape of RBC with both pulling and pushing forces at the dimple and donut regions respectively ($F_{dimple}$ = 3.73 $pN/\mu m^2$ and $F_{donut}$ = 0.53 $pN/\mu m^2$). Adding the small pushing force at the donut region compared to Fig. 4, increased the total shape error noticeably from $\epsilon_{total} \sim 0.22\ \mu m$ to $\epsilon_{total} \sim 0.38\ \mu m$.

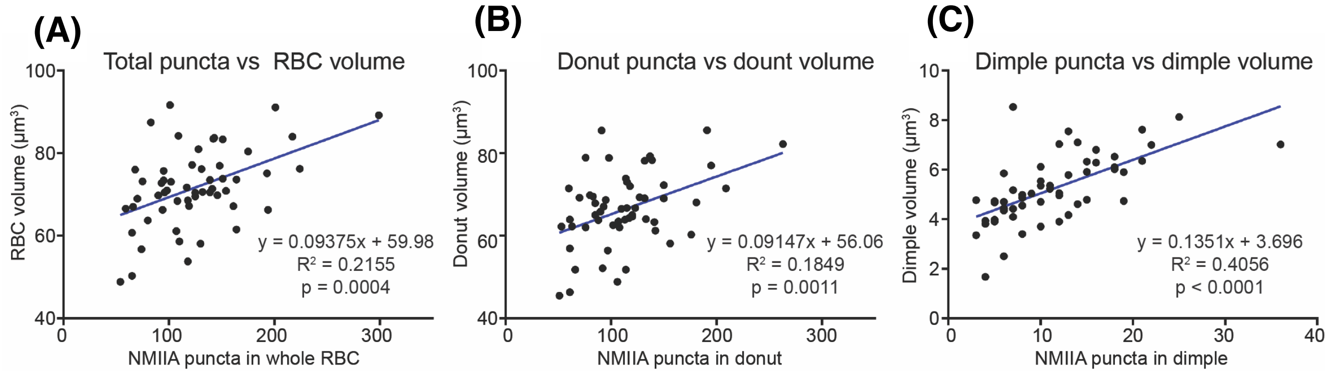

Figure S4: The number of NMIIA puncta increases with region volume. (A-C) Scatterplots of NMIIA puncta in the whole RBC versus the whole RBC volume (A), NMIIA puncta in the donut versus the donut volume (B), and NMIIA puncta in the dimple versus the dimple volume (C). Blue lines represent linear best-fit lines. The equation for the best-fit line, the $R^2$ value, and the p-value for each linear regression are given next to the chart. In all three regions, the number of NMIIA puncta increases with increasing region volume. n = 55 RBCs from 3 individual donors (same RBCs as in Fig. 5).

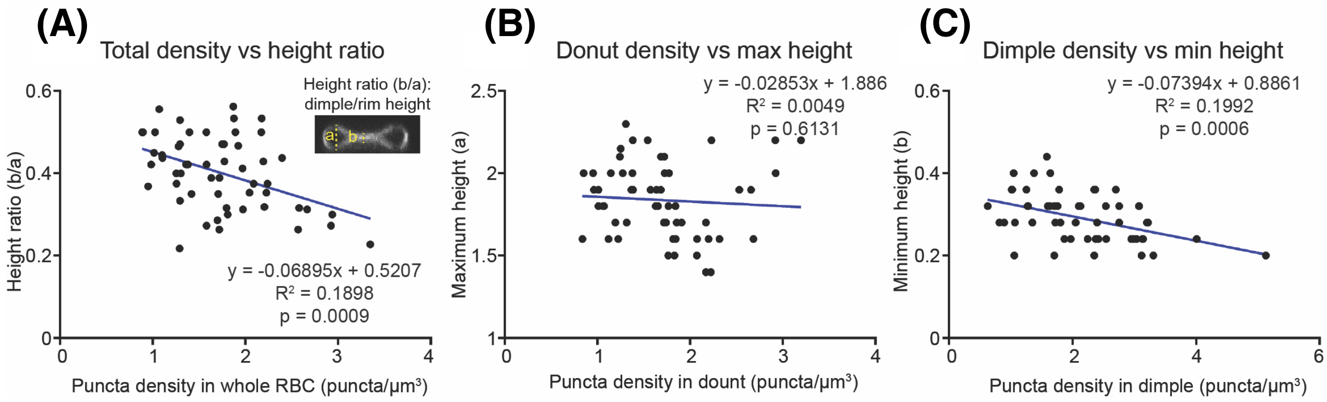

Figure S5: RBCs with higher biconcavity have higher NMIIA density in the whole RBC and in the dimple. (A-C) Predictions about the relationship between NMIIA density and RBC shape based on the model. (A) Scatterplot of NMIIA puncta density in the whole RBC versus RBC biconcavity (height ratio) measured from the ratio of cell height at the dimple to maximum cell height at the rim from an XZ slice near the center of the cell (see inset). (B) Scatterplot of NMIIA puncta density in donut region versus maximum height at the rim (rim height (a) in (A)). (C) Scatterplot of NMIIA puncta density in dimple region versus minimum height at the dimple (dimple height (b) in (A)). (A-C) Blue lines represent linear best-fit lines. The equation for the best-fit line, the $R^2$ value, and the p-value for each linear regression are given next to the chart. Height ratio generally decreases (indicating increasing biconcavity) with increasing whole RBC puncta density (A), and minimum height at the dimple generally decreases with increasing dimple puncta density (C), though there is high variability in this trend between individual cells. Maximum height at the rim does not correlate with donut puncta density (B). n = 55 RBCs from 3 individual donors (same RBCs as in Fig. 5).
